# Ice and air: Visualisation of freeze-thaw embolism and freezing spread in young *L. tulipifera* leaves

**DOI:** 10.1101/2024.10.26.620221

**Authors:** Kate M. Johnson, Muriel Scherer, Dominic Gerber, Robert W. Style, Eric R. Dufresne, Craig R. Brodersen

## Abstract

Spring freezing is an unforgiving stress for young leaves, often leading to death, with consequences for tree productivity and survival. While both the plant water transport system and living tissues are vulnerable to freezing, we do not know whether damage to one or both of these systems causes death in leaves exposed to freezing. Whole saplings of *Liriodendron tulipifera* were exposed to freezing and thawing trajectories designed to mimic natural spring freezes. We monitored the formation of freeze-thaw xylem embolism and damage to photosynthetic tissues to reveal a predictable progression of ice formation across the leaf surface that is strongly influenced by leaf vein architecture, notably the presence or absence of bundle sheath extensions. Our data also show that freeze-thaw embolism occurs only in the largest vein orders where mean vessel diameter exceeds 30µm. With evidence of both freeze-thaw embolism and damage to photosynthetic tissue, we conclude that this dual-mode lethality may be common among other wide-vesseled angiosperm-leaves, potentially playing a role in limiting geographic distributions, and show that bundle sheath extensions may stall or even prevent freezing spread.

**Highlight:** Both ice and air lead likely lead to death in young *L.tulipfera* leaves exposed to freezing, with the spread of both governed by physical characteristics of these leaves.

## Introduction

Freezing is one of the strongest determinants of plant distributions (Burke, Gusta et al. 1976, Loehle 1998, Sakai and Larcher 2012, Muffler, Beierkuhnlein et al. 2016), owing to the unforgiving and often lethal nature of freezing in plant tissues that are not specifically equipped to avoid, resist or compartmentalise ice crystal formation (Burke, Gusta et al. 1976, Stegner, Buchner et al. 2023). Some trees have adaptations to prevent or tolerate freezing within their tissues year-round, while others, such as those in temperate regions, are adapted to seasonal winter-freezing (Burke, Gusta et al. 1976, Wisniewski and Fuller 1999, Sakai and Larcher 2012). Both the xylem and living leaf tissue are critical to consider when assessing the effects of freezing, as damage to plant water transport or the sites of photosynthesis jeopardise tree growth and survival (Brodribb, Brodersen et al. 2021).

Within branches and tree-trunks, xylem anatomy is known to influence the likelihood of freezing damage. During freezing, gas bubbles become suspended in the xylem sap as the crystallization process pushes dissolved gasses out of solution. Upon thawing, these bubbles either dissolve back into solution or expand to fill xylem conduits and block water flow depending on the pressure of the surrounding liquid (Hammel 1967, Sperry and Sullivan 1992, Hacke and Sauter 1996). Species with larger diameter xylem conduits have been shown to be more vulnerable to ‘freeze-thaw embolism’ (Sperry and Sullivan 1992, Feild and Brodribb 2001, Tyree and Zimmermann 2002, Cavender-Bares, Cortes et al. 2005, Sevanto, Holbrook et al. 2012, Li, Luo et al. 2024), which may, in turn, at least partially explain geographic range distributions based on the likelihood of freezing conditions (Sperry 1995). There are a number of possible explanations for this. Larger vessels in stems have been shown to embolise earlier than smaller vessels exposed to freezing (Davis, Sperry et al. 1999, Cavender-Bares, Cortes et al. 2005), and smaller conduits have been shown, in some cases, to freeze at higher velocity which lead to smaller air bubbles (Ewers 1985) which are more easily dissolved upon thawing (Sevanto, Holbrook et al. 2012).

Winter deciduousness allows many angiosperm trees (and some gymnosperms) to avoid leaf-level freezing damage, but does not protect spring growth, nor leaves that persist into autumn. In particular, freezes after bud-burst in spring can have severe consequences for deciduous trees (Burke, Gusta et al. 1976, Vitasse, Bottero et al. 2019), including strong reductions in tree growth (Vitasse, Bottero et al. 2019).

Intracellular ice formation invariably leads to cell death by rupture of the cell membranes, caused by internal nucleation of piercing by external ice crystals (Mazur 1969, Burke, Gusta et al. 1976, Steponkus, Dowgert et al. 1983, Guy 1990). Extracellular ice formation can also result in cell death by creating a water potential gradient which draws out cellular water to feed the freezing-front (Guy 1990, Steponkus and Webb 1992), therefore imposing a desiccation stress resembling that caused by drought (Ruelland, Vaultier et al. 2009, Vitra, Lenz et al. 2017, Yang, Gerber et al. 2024).

Spring frost may be lethal in young leaves through at least two different modes that are not mutually exclusive. First, ice formation in or around the living tissues poses a significant threat because of the potential damage to cell walls and membranes leading to electrolyte leakage, damage to organelles and the generation of reactive oxygen species (Wise 1995, Lütz 2010). Second, embolism formation resulting from the freezing and thawing of the xylem sap could lead to complete failure of the leaf vein network thereby cutting off the supply of water to downstream tissues (Brodribb et al. 2021). While the effects of freeze-thaw embolism have been well documented in stems (Sperry, Donnelly et al. 1988, Feild and Brodribb 2001, Ashworth and Pearce 2002, Cobb, Choat et al. 2007, Mayr, Schmid et al. 2014, Robinson, Rennie et al. 2023), this has never been visualised in leaves.

The primary means of tracking the propagation of freezing damage through whole leaves has been with high temporal resolution using multiple forms of thermal imaging (Larcher, Meindl et al. 1991, Gusta, Wisniewski et al. 2004, Hacker and Neuner 2007, Kokin, Pennar et al. 2018, Zhang, Han et al. 2023). Hacker and Neuner (2007) in particular provide key, novel information about ice nucleation and freezing relative to leaf anatomy, showing ice nucleation beginning in the veins and spreading through the mesophyll tracking leaf venation, however this is one of few papers to consider the influence of leaf venation on ice-spread. While thermal imaging with very high temporal resolution (Hacker and Neuner 2007; 25 images per second) and some optical imaging (Kokin, Pennar et al. 2018) have been used to study leaf freezing; longer-term high resolution imaging of leaves during freezing and thawing has not been conducted. Given that ice crystallization is often lethal in living tissues and that leaf cells are also supplied by a vascular system that is vulnerable to freeze-thaw embolism, one of the goals of the present study was to provide a detailed characterisation of ice formation in leaf tissue by visualising freezing and thawing in the leaves of intact plants using a new application of an existing method, as used in (Kane and McAdam 2024).

Here we visualise the effects of freezing and thawing on both the living cells and the xylem in young leaves in *Liriodendron tulipifera*, a common deciduous tree in North America. To detect possible embolism and track ice crystal formation in leaves we used a combination of time-lapse imaging and chlorophyll fluorescence imaging to non-invasively study young leaves of *L. tulipifera* saplings exposed to freezing conditions, with minimum air-temperatures ranging from −1 to −5 °C. By capturing images of newly expanded leaves at regular intervals throughout freezing and thawing we aimed to elucidate the pattern of freezing spread in these leaves and capture any embolism that may occur using the Optical Vulnerability Method of Brodribb et al. (2017). We then measured chlorophyll fluorescence of leaf tissue before and after freezing-and-thawing as a proxy for tissue damage (Brodribb et al. 2021). To compare the effect of freeze-thaw embolism vs. drought-induced embolism, validating the capacity of the cavicams to detect freeze embolism, and isolating how freezing may differentially impact leaf tissue, we tracked drought-induced embolism in leaves of the same age, sourced form adult trees at various locations.

*L. tulipifera* has been shown to possess bundle sheath extensions (BSEs), structures that connect the bundle sheath and the epidermis and influence a range of physiological process in plants relating to both photosynthesis and hydraulics (Wylie 1943, Wylie 1952, Pray 1954, Esau 1960 ; Fig S1, Zwieniecki, Brodribb et al. 2007). Bundle sheath extensions compartmentalise the leaf lamina restricting the movement of gases therefore allowing gas exchange to occur independently across the leaf surface (Neger 1918), and have been shown to have numerous physiological and biophysical functions (Karabourniotis, Bornman et al. 2000, Zwieniecki, Brodribb et al. 2007, Buckley, Sack et al. 2011, Trifiló, Raimondo et al. 2016, Kawai, Miyoshi et al. 2017). Notably, highly sclerified BSEs have been shown to prevent the lateral prorogation of ice in the lamina in *Cinnamomum camphora*, yet this is the only species in which this has been documented (Hacker and Neuner 2007, Barbosa, Chitwood et al. 2019). In *L. tulipifera* BSEs extend to both the upper and lower epidermises in second order and third order veins (Pray 1954; Fig S1), while they extend only to the abaxial epidermis in higher order veins (Pray 1954, Pieruschka, Schurr et al. 2006, Barbosa, Chitwood et al. 2019). It is possible that even the non-sclerified bundle-sheath extensions in *L. tulipifera* could act as a barrier to the ice crystallization front as it progresses across the lamina, which led us to attempt to determine their influence in this species.

Based on the known physical processes that govern ice nucleation and growth in laminar systems that superficially resemble leaves, we expected ice to first form in leaf regions with low solute content, and therefore a higher freezing temperature, such as the midveins containing numerous xylem conduits loaded with xylem sap. Ice would then propagate outward across the lamina as temperature decreases below the freezing point of the surrounding tissues with higher solute content. Given that the BSEs represent a physical barrier that would slow the movement of water toward the ice nucleation front, we hypothesised that BSEs would significantly influence the spatial pattern and progression of ice across the lamina, which would be most noticeable in the higher order veins where BSEs span both the upper and lower epidermis leading to patchy ice-spread. We developed two possible working hypotheses for freeze-thaw embolism: First, freeze-thaw embolism would occur upon thawing in a pattern resembling embolism caused by drought, where the lower vein orders embolize first, followed by the higher order veins. Second, embolism would occur only in veins with large vessel diameters (equal to or > 30 µm) based on research in stems which shows a steep increase in freeze thaw embolism in conduits above this threshold (Pittermann and Sperry 2006). We predicted that ice crystal formation would lead to damaged cell walls and membranes and thus irreparable damage to the mesophyll, made visible by significant decreases in chlorophyll fluorescence shortly after thawing. If freeze-thaw embolism occurred, it would lead to cell death in any remaining un-frozen tissue but over a longer time period.

## Materials and Methods

This study comprised three core experiments, whereby we first exposed saplings to freezing and thawing cycles while monitoring leaf vein embolism and chlorophyll fluorescence. We then compared those patterns of freeze-thaw embolism to those produced with drought-induced. Finally, we examined the freezing and thawing process at much higher magnification to determine the underlying physical processes.

### Experiment 1: Freezing whole saplings

Saplings of *Liriodendron tulipifera* (∼ 2 years old) were grown under glasshouse conditions (day temperature of ∼25-30 °C and night temperatures of ∼ 15-20 °C) for 1.5 months, until there were at least two fully expanded leaves per tree, before experimentation (conducted between March and May 2022). A total of 13 well-watered trees were frozen to one of five temperature minima (−1, −2, −3, −4 or −5 °C). We selected this range of temperatures to both determine the temperature needed to fully freeze leaves, but also to test the effects of cold temperatures that did not induce freezing. We used the Optical Vulnerability Method (OVT) of Brodribb et al. (2017) to monitor both the formation of embolism but also the changes in optical density of the mesophyll associated with freezing. We imaged a total of 24 leaves, each tree was frozen as described below.

Saplings growing in pots were placed in an upright freezer (Model: FFFH20F2QWC, Electrolux, Stockholm, Sweden) which was attached to a thermostat (Model: ITC-310T-B, Inkbird Tech, Shenzhen, China) which was used to control the freezing trajectory. Two adjacent fully expanded leaves were placed inside ‘cavicam’ imaging clamps (docs.cavicams.com, Fig. 1). Briefly, a digital Raspberry Pi camera (Model: V2, 9132664; Raspberry Pi Ltd., Cambridge, UK) was positioned to observe the adaxial leaf surface and transmitted light is provided by a digitally controlled set of LED lights that only illuminate during image capture (Brodribb, Bienaime et al. 2016). Cameras were set to capture images every 5 minutes during freezing and thawing, a process that was programmed to occur over 20 hours (see details below). These cameras had slightly different magnifications, allowing freezing to be captured at two different scales, but the results were still comparable with no significant difference found in the embolism of percentage of freezing detected between the cameras (ANOVA p> 0.05). A custom-built temperature logger ‘envirologger’ run by a data-logger (Adafruit feather M0 Adalogger; Adafruit Industries LLC; New York, USA) was placed in the freezer to track the internal conditions with a temperature/humidity sensor (Model: AM2315 12C, Adafruit Industries LLC).

**Figure 1:**
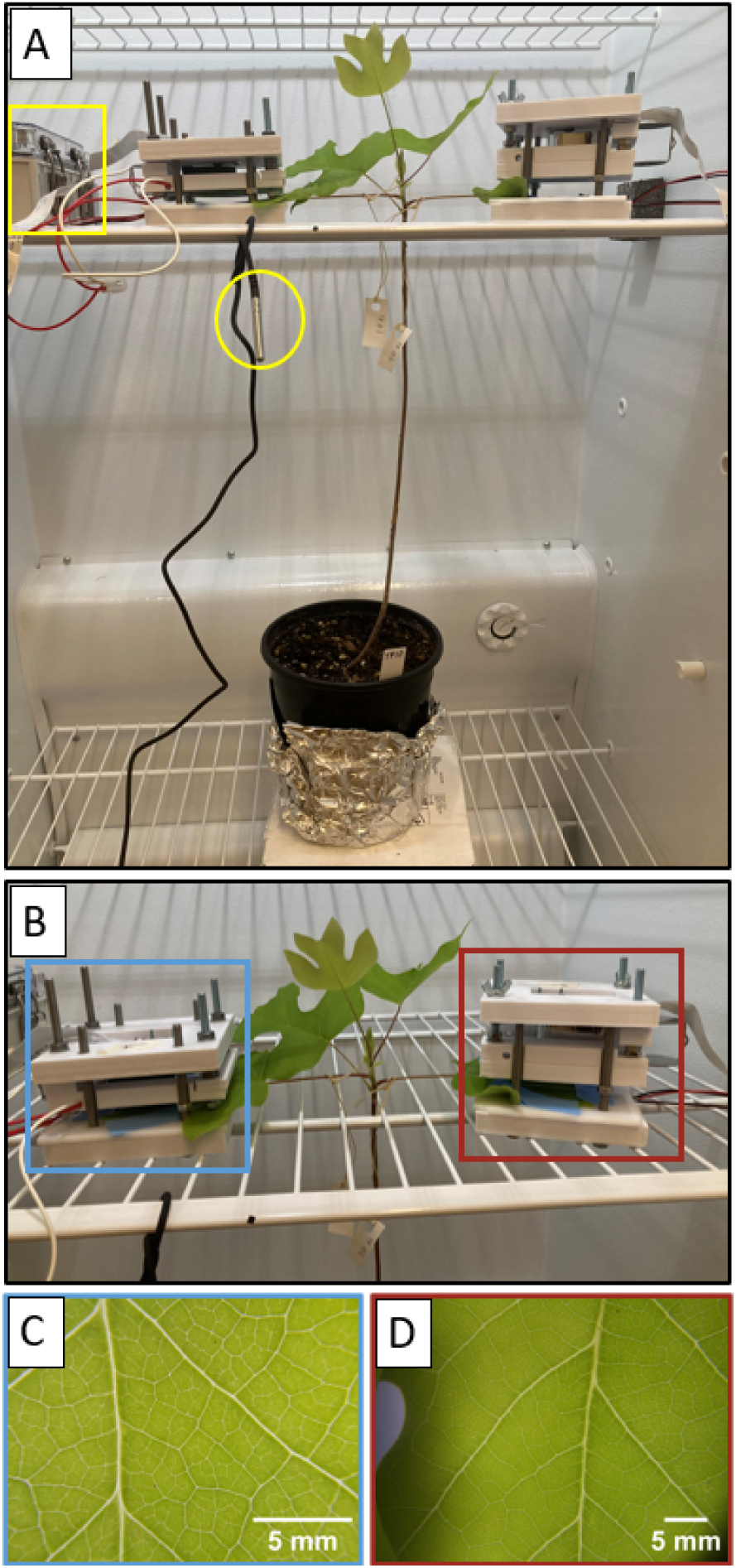
The set-up for monitoring freeze in *L. tulipifera* trees. Panel (A) shows the sapling inside the freezer with cameras attached. The unit for measuring freezer air temperature is denoted by a yellow box while the temperature probe attached to the device controlling the freeze trajectory is circled in yellow. In panel (B) the two cameras are distinguished by boxes, with the higher magnification camera in blue (C) and lower magnification camera in red (D). Corresponding images taken by each camera are shown below the respective cameras, in boxes of the same colours (blue; higher magnification and red; lower magnification) with a 5mm scale bar in the bottom right-hand of each image.

The freezing sequence and trajectory were based on winter/spring temperature data from a weather station (Model: CR300; Campbell Scientific, Utah, USA) monitoring windspeed, air temperature, relative humidity, precipitation, soil moisture and soil temperature, averaged hourly over the measurement period) at nearby Yale Myers Forest (Union, CT, USA; 41° 57’ 8.2944" N, 72° 7’ 26.688" W), where *L. tulipifera* trees are known to grow, in order to mimic the freezing conditions experienced by this species under natural conditions. A representative night in 2019 where temperatures reached −5 °C was chosen as a reference. The temperature data from Yale Myers Forest (recorded half-hourly) was averaged and rounded to the nearest whole number to determine mean hourly temperatures. These temperatures where then used to inform a 12-step programmed freezing sequence. This resulted in a stepped freezing sequence that ran for approximately 20 hours (including an ∼ 6 hour period at a positive temperature at the end) mimicking an overnight freezing event (Fig S2).

### Imaging: Chlorophyll Fluorescence

To determine the physiological effects of freezing temperatures on leaves we used chlorophyll fluorescence imaging, which measures the status of the photosynthetic apparatus and its capacity for processing light. We performed dark-adapted Fv/Fm imaging on leaves in the laboratory both before freezing and thawing to provide a baseline reference measurement, and then again after freezing and thawing to determine how much the photosystems were damaged (Mini Imaging-PAM M-series, Walz GmbH, Germany). Before fluorescence measurements, a small piece of opaque tape was applied to a small section (∼ 0.5x 0.5 cm) of the leaf, then the leaves were wrapped in aluminium foil to exclude light. The patch of opaque tape was used because initial testing showed that, as removal of aluminium foil was required to align the leaf under the imaging PAM, an initial measure was needed to maintain some leaf tissue in the dark-adapted state required for Fv/Fm measurements. The tape was removed seconds before measurement to ensure that the leaf tissue beneath the tape remained fully dark adapted until fluorescence was measured.

### Imaging: Freezing and thawing in the leaf mesophyll

Image sequences were processed in FIJI software (Schindelin, Arganda-Carreras et al. 2012) to determine the spread and extent of ice crystallization during the experiments. The tracing tool in FIJI was used to quantify the total leaf area frozen at each time step, which was indicated by a change in the colour intensity of the pixels. Freezing was detected as patches of brighter pixels in the leaf lamina, typically bounded by higher order veins.

To quantify the colour change in both mesophyll tissue that did and did not freeze, circular Regions of Interest (ROIs) of 40×40 pixels were selected in regions of both mesophyll that froze and that which did not in all leaves frozen to −4 °C. The mean grey scale value in these ROIs was calculated at three time-points, at the start of image capture, at −4 °C (at freezing) and immediately after thawing. The values at −4 °C and after thawing were expressed as a percentage of the initial mean greyscale value (from the start of image capture) and the average percentage of the initial value was calculated for both frozen and unfrozen mesophyll. Mean Fv/Fm for unfrozen, frozen and thawed tissues was calculated from measurements taken across all leaves that froze.

A similar method, utilising circular ROIs in FIJI, was used to track the change in mean greyscale area in the midveins and mesophyll of all leaves which experienced freezing. In each leaf, ROIs which fit within the diameter of the midvein were selected and added to the ROI manager. This was repeated across the image stack or up to 250 images (where more images were recorded), with the ROI shifted using ‘edit>selection>specify’ when the leaf moved. The ‘Interpolate ROI’s’ option through the ROI manager was then used to interpolate between the selected ROIs for all images. The ‘measure’ function was then used to calculate the mean greyscale area in the selected ROI for each image. This was then repeated for a section of mesophyll adjacent to the midvein, using an ROI of the same area. The mean greyscale values across the images were expressed as a percentage of the initial (unfrozen) mean greyscale value in order to generate lines plots of changes in mean grey scale value before, during and after freezing.

The freezing extent was calculated in the image where the maximum amount of tissue was frozen, which corresponded with the minimum air temperature to which the leaves were exposed. The frozen regions were selected using the tracing tool in FIJI (Schindelin, Arganda-Carreras et al. 2012) and expressed as percentage of the total visible leaf area within the field of view to calculate a percentage.

### Imaging: Embolism upon thawing

We monitored the leaf vein network for the formation of embolism using the OVT (Brodribb, Bienaime et al. 2016). Embolism was monitored by the cameras for 12-24 hours post thawing to ensure that all embolism was captured. Freeze-thaw embolism was analysed using the same methodology which is used to highlight drought-induced embolism with leaf images analysed via an image subtraction, where each image is subtracted from the one before to reveal the changes between the images. As there is a change in light transmission as air rapidly replaces water in the xylem due to embolism, the image subtraction reveals these changes which can then be visualised through time. A detailed description of the methodology used to process images can be found here: https://www.opensourceov.org/. While embolism was expected to occur upon thawing, cameras remained attached to plant organs for ∼6 hours to ensure that all embolism was captured.

### Experiment 2: Drought embolism in additional leaves

Optical analysis of drought-induced embolism was undertaken in a total of six leaves from branches of adult trees, one leaf was sourced from a tree located on the Yale University campus, New Haven, Connecticut, USA (41°19’16.6"N 72°55’25.8"W), and the remaining five leaves were sampled form across four trees located in North Hobart, Tasmania, Australia (42°52’07.6"S 147°18’57.8"E). Young, fully expanded leaves (as a similar developmental stage as those used in the freezing experiment) were chosen for imaging. One leaf from each branch was placed within an imaging clamp and set to take images at 5- minute intervals as the branches dried. Embolism was determined to have finished after 12 hours without embolism. Images were analysed using an image subtraction, as described above.

### Analysis: Embolism in Experiment 1 and Experiment 2

The timing and extent of embolism were determined both in leaves exposed to freezing and those exposed to drought. To visualise embolism through time, embolism was colour coded according to time in both a leaf exposed to freeze and a leaf exposed to drought. This was done using FIJI (Schindelin, Arganda-Carreras et al. 2012) using the ‘colour slices’ function in the ‘OSOV toolbox’, the toolbox and instructions are available here: https://www.opensourceov.org/.

### Analysis: Embolism and vein order

To determine how much the vein network embolised, the total length of the visible venation network was calculated along with the total area of the embolised vein network using the ‘segmented line’ tool in image J and the embolised vein area was then expressed as a percentage of the vein network. Total vein area was calculated from a raw image of each leaf taken from the stack of images which captured during the drought or freezing treatment while total embolised vein area was calculated form the ‘Z- stack’ showing all the embolism which occurred during the treatment. Total embolised vein area was then expressed as a percentage of total vein area visible with the optical cameras. As embolism was only observed in the mid-rib and major veins in response to freeze, only these vein orders were included in calculations of vein length and embolised vein length for frozen leaves. The raw images of the vein network and Z-stacks of embolism from leaves exposed to drought were segmented into four even quadrants using a macro in FIJI and the top right corner was analysed for each leaf in order to provide a random sample of the vein network.

In *L.tulipifera* leaves, the midvein, second order and tertiary veins are deemed the ‘major veins’ while the fourth, fifth and sixth and seventh order veins are considered the ‘minor’ veins and make up the most of the total vein length (Pray 1954).

To quantify the contribution of the midvein and second order veins to the earliest embolism is leaves exposed to drought, 25% (embolised area) of the total drought-induced embolism was chosen as a reference value. This value was used to determine whether embolism observed due to freezing and thawing matched early embolism during drought. At this percentage embolism, in all droughted leaves, we calculated both the area contribution of these veins compared to higher vein orders, and the total length of midvein and second order veins embolised as a percentage of the total embolised area of these veins at 100% embolism. For area, the ‘freehand sections’ tool in FIJI was used to ‘circle’ all vein orders expect for the midvein and second order veins and the area of these veins was calculated using the ‘measure’ function and expressed as a percentage of the total cumulative embolised area at 25% embolism. The resulting percentage of higher-order-veins was subtracted from 100% to reveal the percentage contribution of the midvein and second order veins to the embolised area at 25 % embolism. For vein length, the ‘segmented line’ tool was used to calculate the length of the total embolised area of the midvein and second order veins in a Z-stack of the cumulative embolism at 25% embolism. This was expressed as a percentage total embolised midvein and second order vein length at 100% embolism.

### Experiment 3: High resolution imaging of freezing

Branches of adult *L. tulipifera* trees were excised from individuals on the EPFL campus, Lausanne, Switzerland (46.5191° N, 6.5668° E) and transported to the ETH Honggerberg Campus in Zurich. Small leaves were mounted between two glass microscope slides and placed on a customised freezing stage beneath a compound microscope using confocal microscopy (Yokogawa Spinning Disk and Nikon Eclipse, set up &software by 3i imaging). The set-up used is described in (Gerber, Wilen et al. 2023) and the documentation is available at: https://github.com/dogerber/temperature_gradient_microscopy_stage/tree/main. A CF640 filter was used to visualise eaves, capturing the autofluorescence of chlorophyll therefore highlighting the cells in the images.

### Anatomical assessment: Xylem conduit diameter analysis

Young *L. tulipifera* leaves were collected from trees and brought to the lab. Vein samples from three leaves were used for the conduit diameter analysis. Midvein samples were collected 1cm from the petiole lamina junction. Second and third order veins were collected 0.5 cm from the junction with the subtending lower order vein. Samples were mounted to a freezing stage (BFS-3MP, Physitemp Instruments, USA) on a sliding microtome encased in a dilute glucose solution. Thin sections were stained with a 0.5% toludine blue solution, rinsed in DI water, and then mounted in water. Images were captured with a digital camera mounted to a compound light microscope (BX43, Olympus Inc, Japan) using a 4x or 10x objective lens. Images were then used to measure xylem vessel diameter in FIJI. Vessel diameter was measured as the longest distance across the lumen.

### Data analysis

Data were plotted and analysed using SIGMAPLOT v.12.5 (Systat Software Inc, San Jose, California) and RStudio was used to perform Single factor ANOVAs to determine statistical differences between groups in RStudio (2016). Xylem vessel diameter analysis was performed in RStudio using the dplyr package (Yarberry and Yarberry 2021).

## Results

### Pattern of Freezing and thawing

We observed mesophyll freezing in all leaves which were exposed to minimum air temperatures of −4° C or −5° (total of 18 leaves). At −4° C greyscale pixel values increased in the freezing regions that can be interpreted as a change from air to water in the intercellular space as water is drawn out of the mesophyll cells to feed the freezing front (Fig. 2).

**Figure 2:**
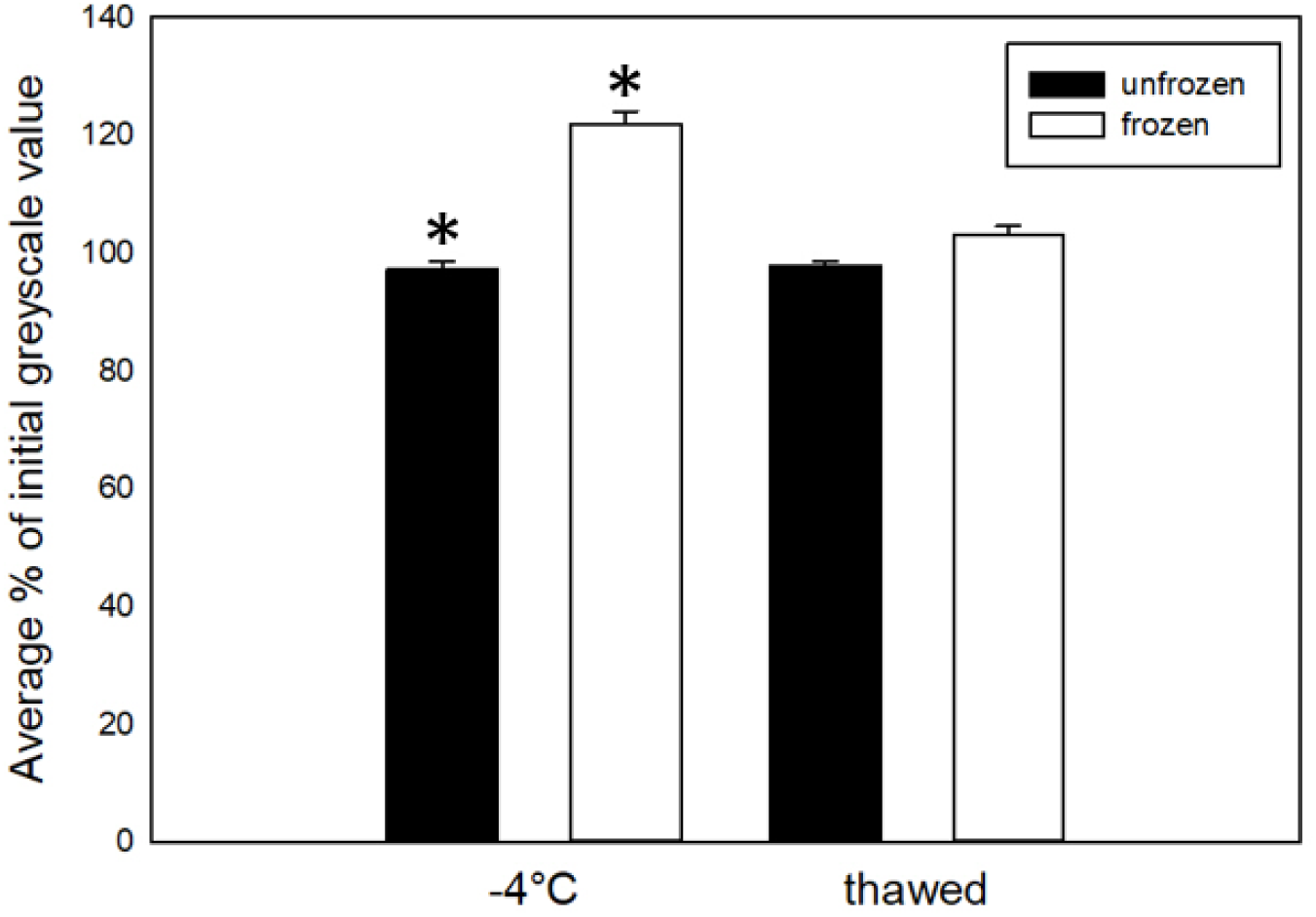
The average percentage of initial (pre-treatment) mean greyscale value (brightness) for all 18 leaves that froze, comparing tissue that did and did not freeze within the same leaves. These values were calculated at −4 °C and immediately after thawing. Unfrozen tissue is shown in black, while frozen tissue in shown in white. This was calculated for all 18 leaves that froze. Asterix’s indicate a statistically significant difference (ANOVA, P<0.05). As these values were calculated as a percentage of the mean greyscale value at the first image in the image sequence, some values were < 100% due to small changes in lighting during image capture.

While an increase in mean greyscale value, indicating freezing, was always detected in the mesophyll, this was not consistently observed in the midvein (Fig 3). Freezing in the mesophyll resulted in a 10-20% increase in mean greyscale brightness while freezing in the veins incurred an increase in mean greyscale value of <5% (Fig 3). Only one of the eight leaves frozen to −4°C showed a signal of freezing in the midvein, while six of the ten leaves frozen to −5°C showed this signal of freezing.

**Figure 3:**
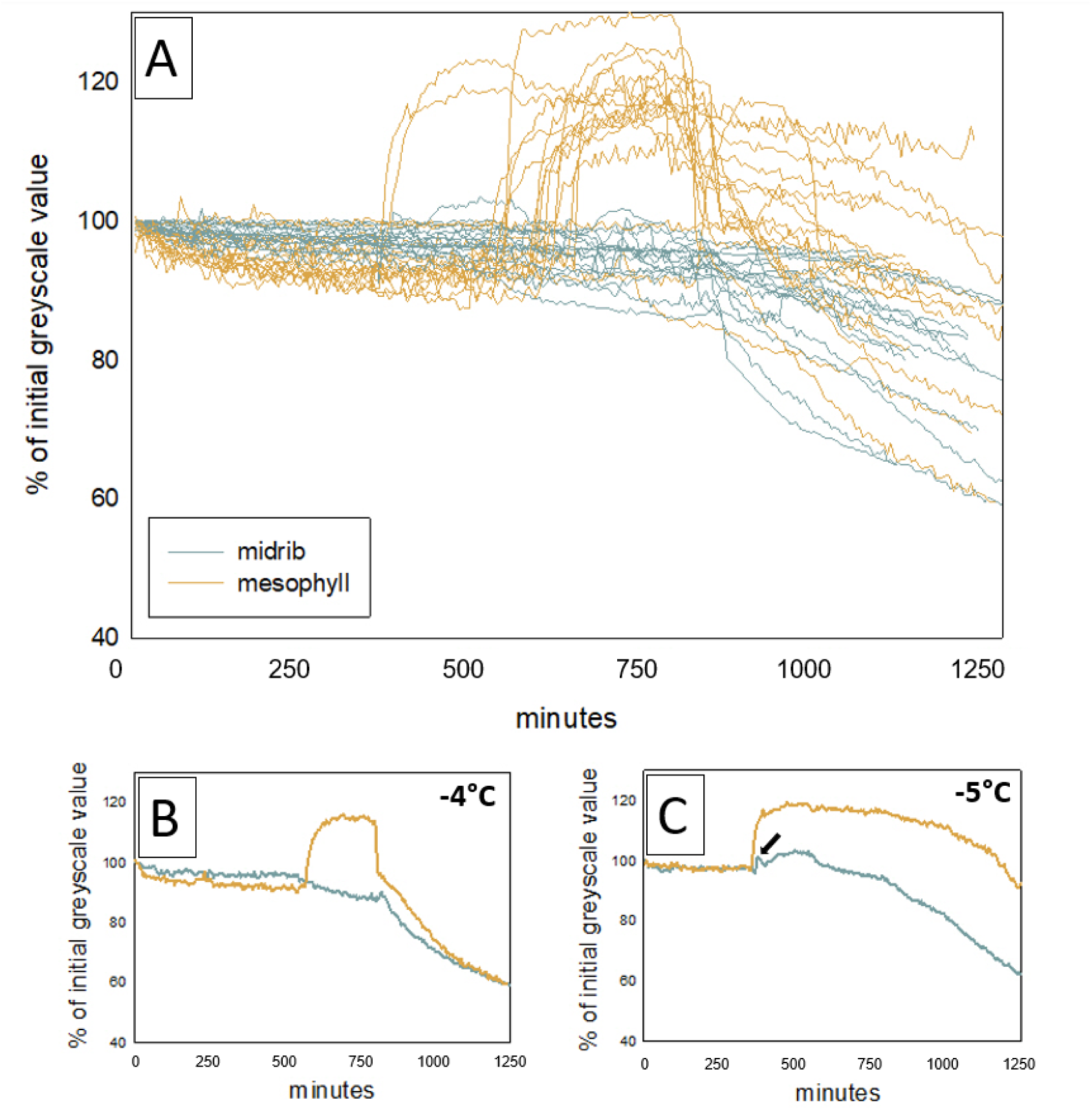
Percentage of initial greyscale value over time for the midvein and mesophyll for 18 *L. tulipifera* leaves frozen to −4°C or −5°C. The mean greyscale values for leaf midveins (blue) and mesophyll adjacent to the midvein (orange) across 250 images (∼20 hours) for all leaves frozen to −4°C or −5°C (A). Increases in mean greyscale brightness in the mesophyll (orange) indicate mesophyll freezing. Panel (B) shows the midvein and mesophyll mean greyscale traces for a representative single leaf frozen to −4°C and panel (C) shows the same for a leaf frozen to −5°C with a black arrow denoting the increase in mean greyscale brightness indicating freezing in the midvein. Panels (B) and (C) serve to highlight two examples and do not describe the pattern observed in all leaves frozen to −4°C and −5°C respectively. The signal and timing of mean greyscale increases varied across of leave independent of temperature.

Freezing was first detected in mesophyll immediately adjacent to the midvein and 2° veins, followed by ice propagation away from the vein into areoles bounded by higher order vines (3-5° veins) (Fig. 4). Freezing began at ∼ −4°C continuing until 0°C when thawing occurred (Fig. 4). Freezing advanced across the leaf surface in a stepwise pattern between adjacent areoles in discrete patches. Freezing was in some cases not perfectly spatially homogenous, and in all but two leaves where 100% of the area frozen (16 out of 18 leaves) some areoles bounded by third order veins remained unfrozen (Fig. 4, Vid. 1, Fig. S3).

**Figure 4:**
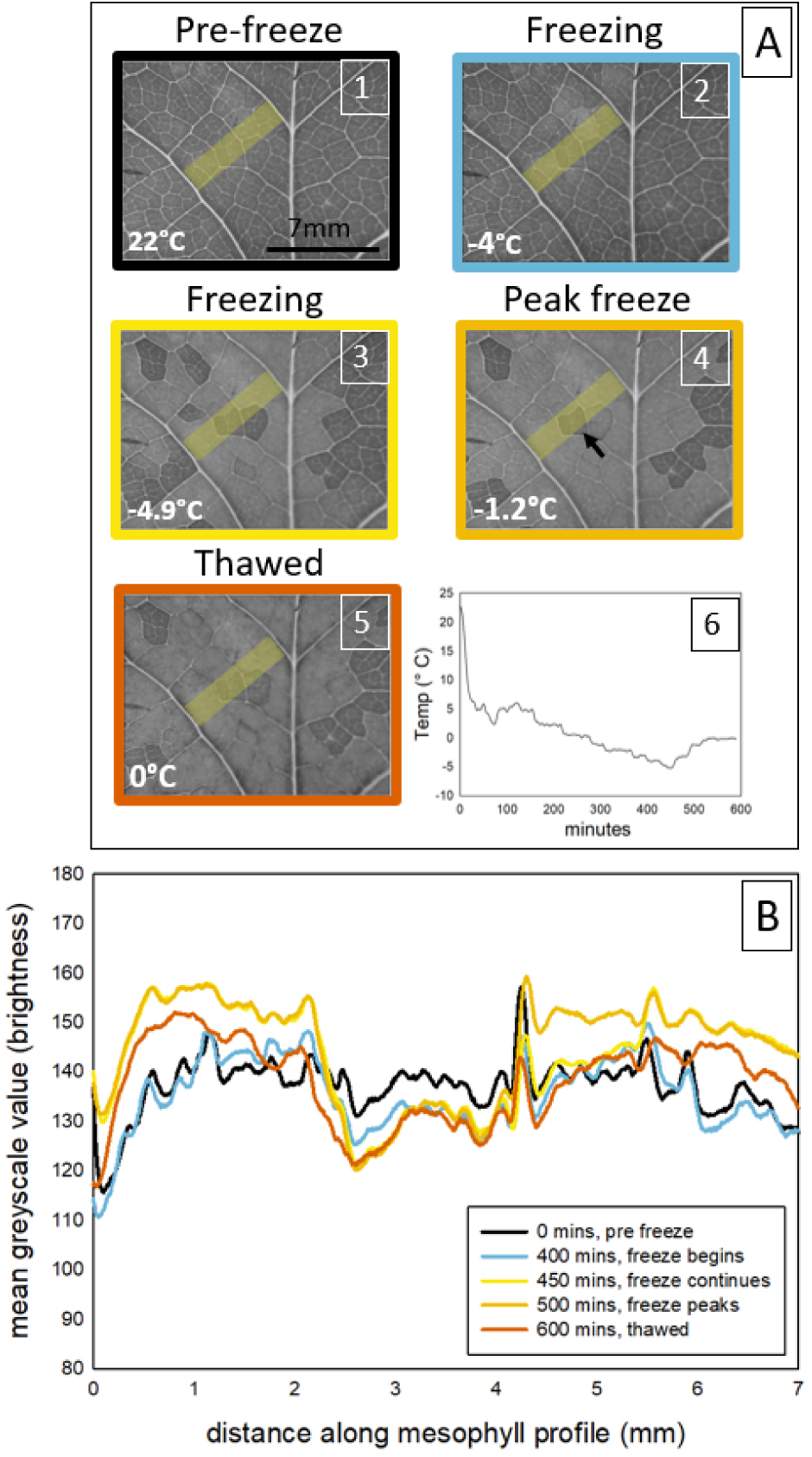
Representative freezing and thawing sequence including images, mean greyscale value and air temperature for a representative *L. tulipifera* leaf visualised using time-lapse imaging. Panel (A) shows greyscale images of the leaf surface including the midvein and surrounding higher order veins with the transect used to calculates mean greyscale values denotes by a yellow rectangle. Mean greyscale values (B) across a 7mm mean greyscale profile between the midvein and a major vein of *L. tulipifera* leaves measured at 5 stages of freezing in a leaf frozen to −5°C [pre-freeze (1): black, freezing; blue (2) and yellow (3), peak freeze: light orange (4), thawed; dark orange (5)]. The temperature at each stage is in the bottom right-hand corner of each image and panel (6) shows time (minutes) vs temperature (°C). This figure shows the distinctive colour change which occurs with freezing visually (A) and numerically (B), illustrating a situation where a patch in the centre of the mean greyscale profile remains unfrozen (denoted by a black arrow in the ‘Peak freeze’ image in panel (B).

Freezing at high magnification in young leaves from adult trees revealed a similar pattern, where a freezing-front propagated through individual areoles, but then stalled upon reaching a third-order vein boundary (Video 2). The freezing front can be observed progressing from the bottom left corner towards the middle of the field of view where it can be seen moving through cells before reaching a third-order vein boundary and then progressing to the bottom right and moving towards the middle before reaching the vein boundary in the same manner (Video 1).

Thawing, by contrast, did not appear to occur as a staged process, occurring over 5-10 minutes images when the temperature rose above 0 °C, Fig. 4). Thawing resulted in a ‘darkening’ of the previously lighter frozen patches under the optical cameras which used transmitted light (Fig. 2, Fig. 4), while thawing corresponded with lightening in the high-resolution video of thawing as this set-up utilised reflected light (Video 3). In the hours following thawing, previously frozen tissue gained a red colouration (Fig. 5).

**Figure 5:**
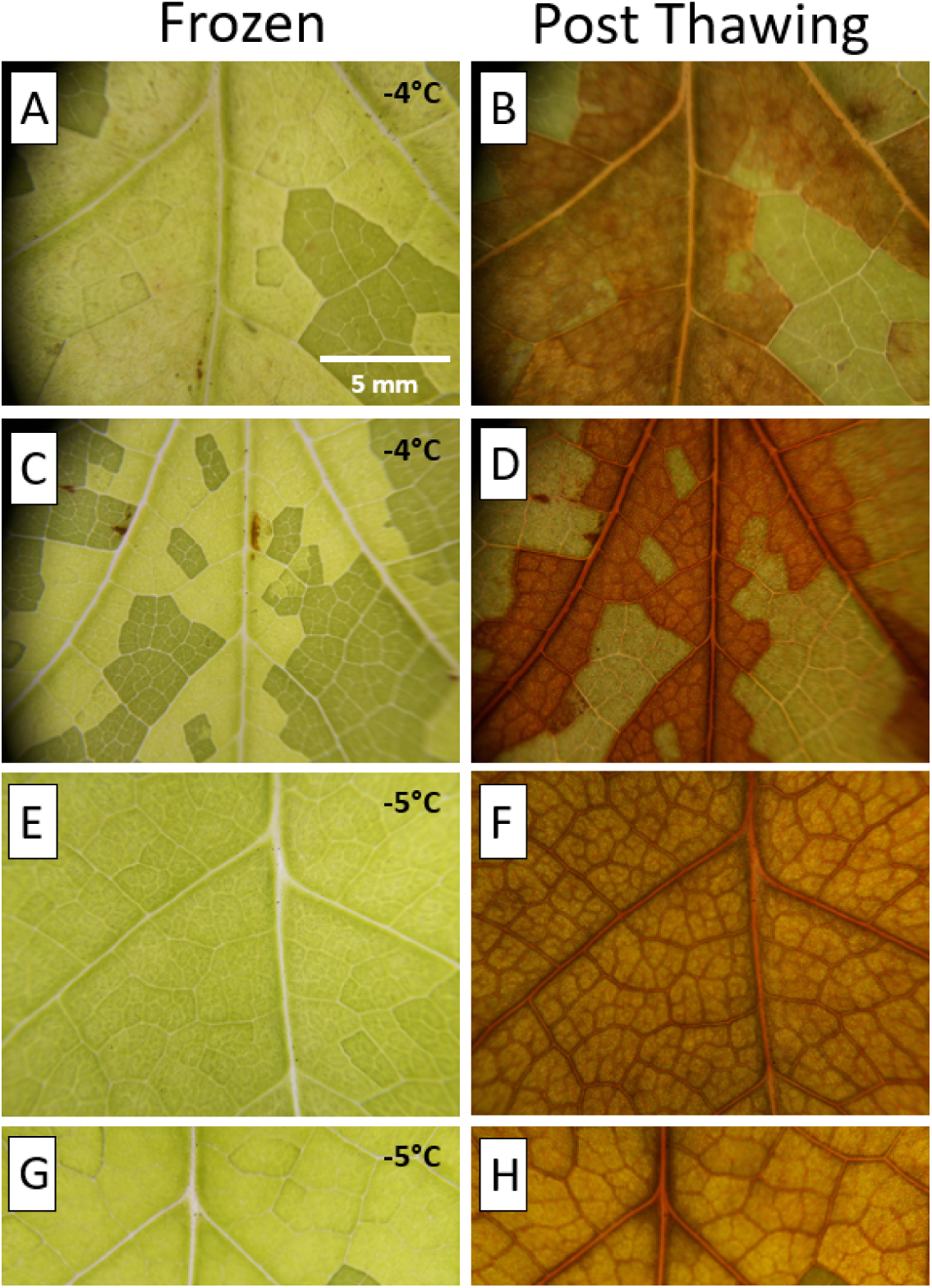
L. *tulipifera* leaves (n=8) at their maximum freezing extent (left hand column) and ∼ 10 hours after thawing (right hand column), demonstrating the reddening of the tissue that occurred in frozen tissue. Image pairs (A, B) and (C, D) show freezing and reddening I leaves frozen to an air temperature of −4° C, while (E, F) and (G, H) show this in leaves frozen to −5 ° C (c). The 5mm scale bar in (a) is true for all panels.

### Freezing extent

Freezing began at air temperatures below −4 °C, with no visible evidence of freezing in leaves exposed to air temperatures of −1°C, −2°C and −3°C. In leaves exposed to air temperatures below −4°C (n = 8), an average of 58%± 5% of the leaf froze within the camera’s field of view. The mean percentage of the leaf turning red increased to 93% ±2% in the ten leaves frozen to −5°C (; P< 0.000008, ANOVA; Fig. 6).

**Figure 6:**
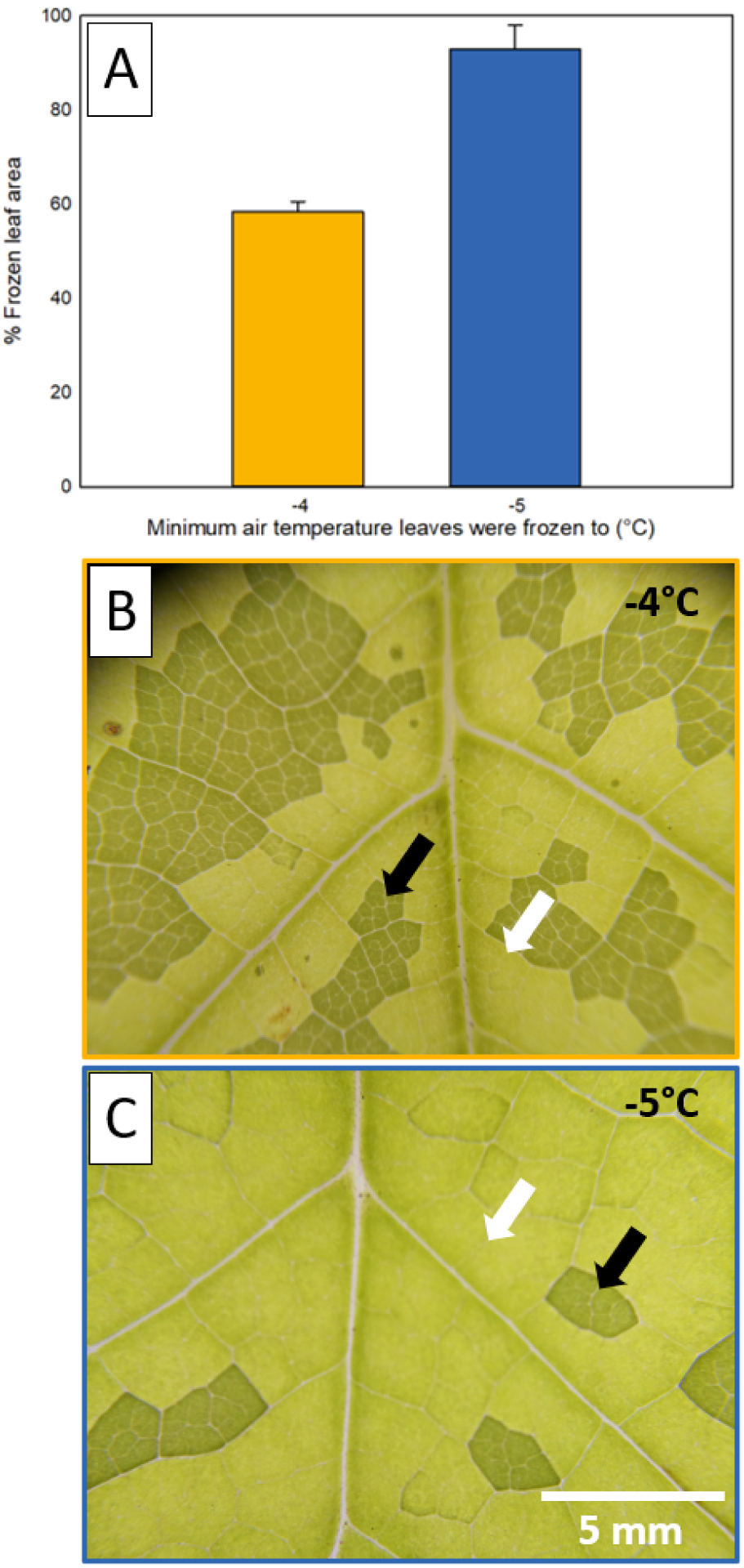
The percentage frozen leaf area in leaves frozen to air temperatures of −4°C and - 5°C. panel (A) shows to total percentage frozen leaf tissue in all leaves frozen to −4°C (gold) and −5°C (blue). Representative images of the freezing extent are also shown in both a leaf frozen to −4°C (B) and a leaf frozen to −5°C (C). Arrows mark unfrozen tissue (black arrows) and frozen tissue (white arrows) in both (B) and (C). The 5mm scale bar in (C) is true for (B) and (C) panels.

### Chlorophyll Fluorescence Imaging

Chlorophyll fluorescence measured ∼6 hours after freezing and thawing invariably decreased in frozen tissue and largely remained high (close to 0.8 Fv/Fm) within leaves that did not freeze and in unfrozen tissue within leaves that did freeze (Fig. 7). Mean Fv/Fm for frozen tissue (0.349 ± 0.06) was significantly lower than that for unfrozen tissue (0.733 ± 0.21; P< 0.000003, ANOVA). In leaves exposed to −1°C −2°C or −3°C, where we observed no freezing of the lamina, fluorescence remained > 0.7 Fv/Fm (Fig S4). Evidence of both Fv/Fm depression and high Fv/Fm values in frozen and unfrozen tissue, respectively, were measured in leaves exposed to temperatures of −4°C or −5°C. The ‘patchy’ nature of freezing in the leaf mesophyll visible with the optical camera (Fig.6) was also visible through chlorophyll fluorescence imaging (Figs. 7b, S5). A lack of fluorescence decline in leaves where freezing was not observed, (exposed to −1°C, −2°C and −3°C air temperatures) indicates that fluorescence values were not impacted by the time spent in the freezer or the placement of cameras on leaves (Fig. S4).

**Figure 7:**
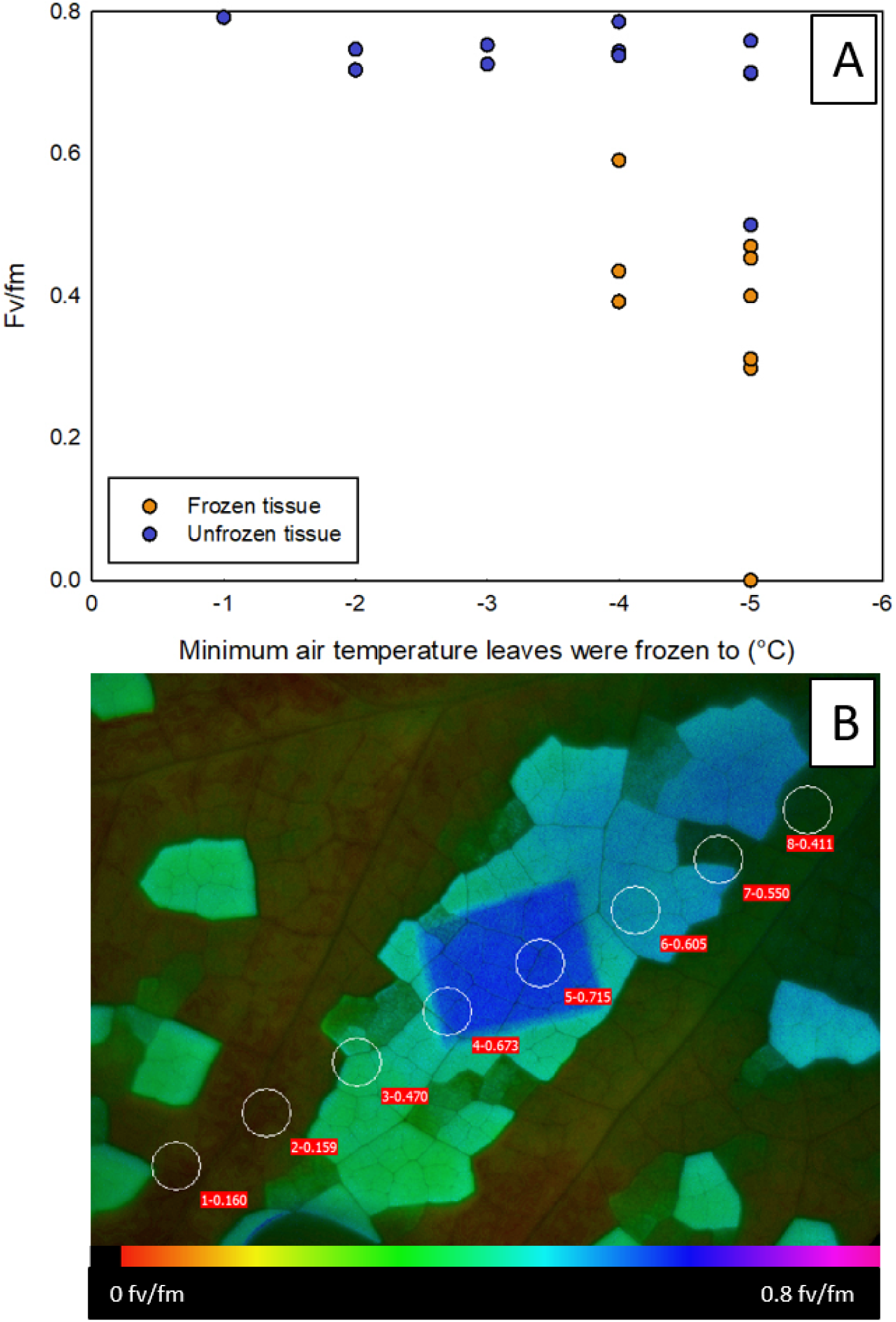
Changes in chlorophyll fluorescence in response to negative temperatures in 24 leaves of *L. tulipifera*. Panel (A) shows Fv/Fm (photosynthetic capacity) in frozen (orange) and unfrozen (blue) leave tissue against the minimum air temperature leaves were frozen to, along with a visual representation of fluorescence in a leaf post-freeze (B).The dark blue rectangle in (B) shows the patch of unfrozen tissue which was ‘dark-adapted’ before the fluorescence measurement, with a value of 0.715 Fv/Fm. Unfrozen tissues outside of this dark-adapted area are turquoise and light green, indicating a higher fluorescence than the surrounding, darker unfrozen tissue. Point values of Fv/fm are shown in eight circular areas of interest arranged diagonally across the image. The colour-scale beneath panel (B) indicates fluorescence values.

### Tree Recovery

While leaves exposed to air temperatures of −4°C or −5°C invariably died, all but one of the 13 trees exposed to these temperatures resprouted leaves (Fig. S7). Resprouting occurred from dormant buds below the leaves frozen during the experiment (Fig. S7).

### Freeze-thaw vs. drought-induced embolism

Freeze-thaw embolism was seen in six of the eight leaves frozen to −4°C, but not in any of the leaves frozen to −5°C. Embolism after freezing occurred in 1-3 events and were isolated to the midvein and 2° veins (Fig. S6; Fig.8). We observed no freeze-thaw embolism in the higher order veins, which was distinctly different than the pattern of embolism spread resulting from drought (Fig. 8). Importantly, the clear visibility of embolism in higher order veins in leaves exposed to drought means that the lack of freeze-thaw embolism in higher order veins was not a failure of the optical cameras to detect these events.

**Figure 8:**
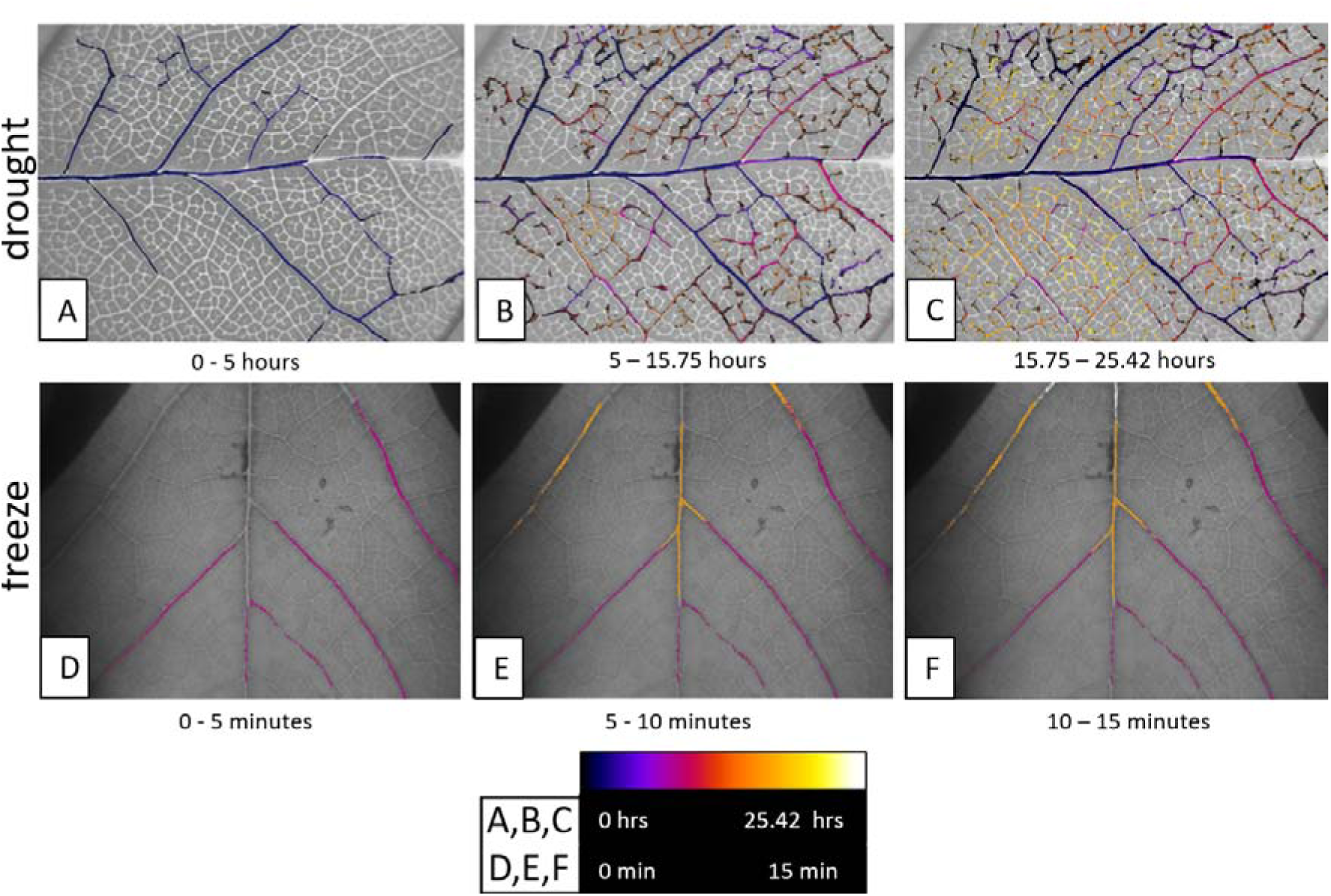
Representative image sequences showing the difference between drought- and freeze-thaw-induced embolism in *L. tulipifera* leaves. Drought-induced embolism spread was detected in young leaves of adult trees and occurred over approximately 1 day, 25.42 hours (A, B, C). Freeze-thaw embolism detected in whole saplings occurred over 15 minutes (D,E,F) in *L. tulipifera* leaves. Panels a, b and c show the progression of drought-induced embolism in four steps of ∼23.5 hours each, while the three images in panel (B) show the progression of freeze-thaw embolism in three steps of 5 minutes, where thawing began at 0 minutes. Embolisms are colour coded according to when they occurred, as indicated by the colour scale beneath the panels.

Freeze-thaw embolism was confined to the midvein and second order veins, while drought induced embolism was observed in all vein orders (Fig. 8). In leaves where freeze-thaw embolism was observed, 77.6 ± 11.8 % of the midvein and second order veins embolised, compared to 100% of these veins in drought-exposed leaves. In response to drought, the percentage of embolism in tertiary veins and above was a similar to the percentage of freeze-thaw embolised midveins and major veins 74.25 ± 7 %, including all vein orders in drought-exposed leaves (adding major and second order veins) did not alter the percentage of the vein network that was embolised substantially, with 78%± 1.4% of the total vein network found to embolise in leaves exposed to drought. The midvein and second order veins often embolised first in response to drought embolism (representing 83.56% ± 4% of the total embolised area at P_25_ and 76% ± 34% of the total embolised length of midveins and second order veins at P_25_ as a percentage of the total length at P_100_). However, embolism in these veins was accompanied, or shortly followed, by embolism in the minor veins as well (Fig. 8) unlike in leaves exposed to freezing and thawing. The timescale across which drought embolism and freeze thaw-embolised were also very different. While freeze-thaw embolism events occurred within 5- 15 minutes, drought induced embolism occurred over a range of 1- 5 days (Fig. 8).

### Xylem vessel diameter

We measured vessel lumen diameter distributions in the three lowest vein orders to determine whether this trait might influence our observations of embolism in this species given that previous work indicates that freeze-thaw embolism vulnerability increases when mean conduit diameter exceeds 30µm (Pittermann and Sperry 2003). We found a mean vessel diameter of 34.4± 6.96, 27.4± 6.95, and 7.45 ±1.85 for the midveins, second order veins, and third order veins, respectively (Fig. 9). We found that 72% and 33.3% of the vessels were greater than 30µm in the midveins and second order veins, respectively, but no vessels >30µm were found in high order veins.

**Figure 9:**
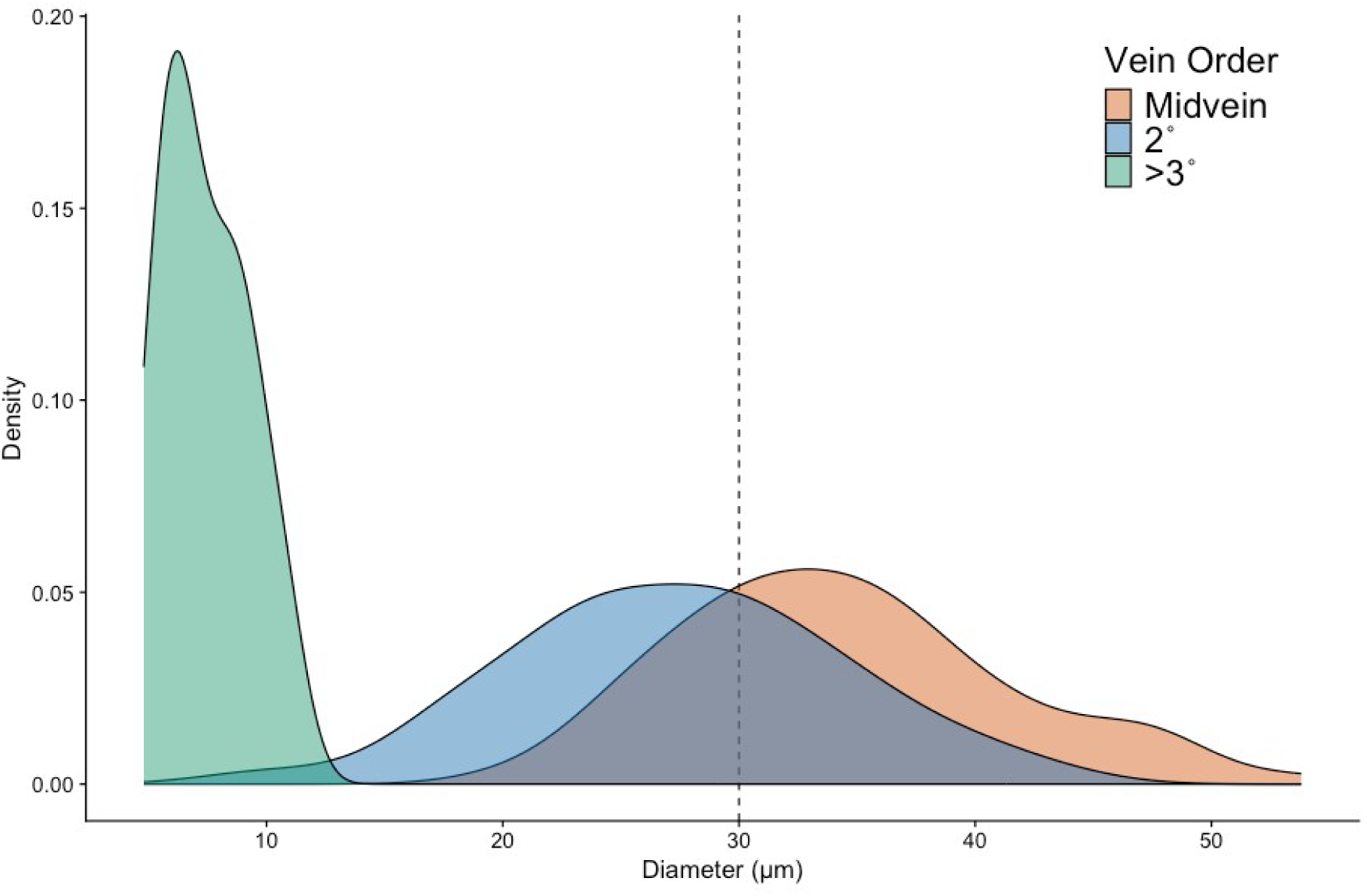
Xylem vessel lumen diameter distributions for leaves of *L. tulipifera*. Distributions are shows for different vein orders (blue = midveins, green = second order veins, orange = high order veins). n = 3, mean diameter is statistically different for each group (ANOVA, p<0.05). Vertical dashed line shows the 30µm threshold, where conduits below this threshold are known to be less vulnerable to freeze-thaw embolism.

## Discussion

Freezing young *L. tulipifera* trees to air temperatures of −4 °C or −5 °C invariably led to visible freezing in the mesophyll, cell damage after thawing, and ultimately leaf death. Our visualisations of ice formation revealed that mesophyll freezing began adjacent to the midveins and second order veins before progressing in a step-wise pattern across areoles bounded by third order veins. These freezing patterns support our hypothesis that bundle-sheath extensions (BSEs) strongly influence how ice propagates and spreads through the lamina of *L.tulipifera* leaves. Freeze-thaw embolism occurred only in veins containing vessels with diameters 30 µm or higher, in contrast to drought-induced embolism which occurred across all vein orders. This finding supports our second hypothesis that freeze-thaw embolism is driven by conduit size, generally agreeing with research in stems (Pittermann and Sperry 2003), providing further support that conduit diameter is a major driver of freeze-thaw embolism across multiple plant organs.

### Mesophyll freezing and thawing

Mesophyll freezing was associated with an increase in tissue brightness, which was also recently shown by Kane and McAdam (2024). This increase in brightness can be explained by the transition in optical density of the intercellular airspaces as the air is replaced with liquid and then solid water during the freezing process. This is the same principle that underpins the Optical Vulnerability Technique (OVT) for the observation of drought-induced embolism (Brodribb, Bienaime et al. 2016, Brodribb, Carriqui et al. 2017). The OVT detects embolism in leaves as a colour-change from light-coloured translucent xylem (water-filled) to darker air-filled xylem, due to the change in optical density of the conduits (Brodribb, Carriqui et al. 2017). Therefore, the transition from darker to lighter tissue at freezing temperatures (Kane and McAdam 2024) can likely be explained by the movement of water/ ice into previously air-filled spaces. As the refractive indexes of liquid water (1.33) and ice (1.31) are very similar, the flooded intercellular space would appear brighter even after thawing, as we saw here (Fig. 4). This colour change was reversed in the high-resolution freezing, whereby freezing caused a darkening of tissue (Video 2). The occurrence of intercellular freezing is in agreement with the idea of freezing as a dehydration stress in plants (Guy 1990, Steponkus and Webb 1992, Ruelland, Vaultier et al. 2009, Vitra, Lenz et al. 2017, Yang, Gerber et al. 2024) whereby water is pulled out of living cells to feed the freezing front in the spaces between cells. In the absence of intracellular freezing, this process is thought to cause tissue death (Steponkus and Webb 1992).

We attribute the difference in the wave-like spread of freezing from around the midvein and second order veins and the stepwise, ‘patchy’ freezing that occurred in mesophyll bound by tertiary veins (Videos 1, 2) to the presence or absence of bundle-sheath extensions (BSEs). previously published work shows that freezing initiated close to the midvein and second order veins (Hacker and Neuner 2007). BSEs extend to both the upper and lower epidermis in tertiary veins of *L. tulipifera* (Fig. S1), but are not present in the midvein or second order veins. BSEs have been shown to be discontinuous or extend to only one leaf surface in fourth order veins and higher in *L. tulipifera* (Pray 1954). The absence of BSEs in the two lowest vein orders likely accounts for the apparently uninhibited spread of ice adjacent to the midvein and second order veins while their presence may explain the independent and patchy freezing or areoles bounded by tertiary veins containing BSEs (Fig S3). The presence or absence of BSEs can therefore be used to explain the delay to freezing spread at tertiary vein boundaries (Videos 1, 2), or complete blockage in others (Figs 4, 6, 7), where BSEs are acting as physical barriers to the movement of the ice crystallisation.

### Freeze-thaw xylem embolism

The most convincing driver of freeze-thaw embolism here may be xylem conduit diameter. Freeze-thaw embolism in the lamina was confined to the two lowest vein orders (midvein and second order veins), which also possess the largest vessels (Figs. 8, 9). We found vessel lumen diameters exceeding 30µm in both vein orders showing embolism. In contrast, we observed no embolism in veins with vessels <15µm in diameter veins. This observation aligns with research in conifer stems which shows a sharp increase in vulnerability to freeze-thaw embolism when conduit diameter exceeds 30µm (Pittermann and Sperry 2003) and is consistent with the large body of research in woody plants which shows that larger diameter conduits are at greater risk (Sperry and Sullivan 1992, Sperry, Nichols et al. 1994, Feild and Brodribb 2001, Tyree and Zimmermann 2002, Pittermann and Sperry 2006, Choat, Medek et al. 2011, Robinson, Rennie et al. 2023). This has been theorised (Sevanto, Holbrook et al. 2012) and been inferred to be the case in leaves (Ball, Canny et al. 2006) but has, until now, not been visualised.

A reduction in xylem conduit diameter was one of the key adaptations that allowed flowering trees to radiate into freezing environments (Zanne, Tank et al. 2014) and decreasing conduit diameter with increasing probability of freezing temperatures has been observed at a global scale (Sperry 1995, Sevanto, Holbrook et al. 2012). With a large population of vessels <30µm, it is possible that if embolism of the lower vein orders was incomplete, the smaller conduits which are theoretically less vulnerable to freeze-thaw embolism could maintain water supply to these leaves. This may allow *L. tulipifera* leaves to resist freezing temperatures provided that photosynthetic tissues are ‘hardened’ to freezing temperatures. The extent of freezing damage in photosynthetic tissues may, however, outweigh any effects of embolism, at least early in the growing season when leaves are not acclimated for cold temperatures.

A mean greyscale signal of freezing in the midvein was observed in six out of the 10 leaves frozen to −5°C but only one out of the eight frozen to −4°C. This is contrary to evidence from thermal imaging which showed that freezing occurs first in the veins (Hacker and Neuner 2007). It is possible that vein freezing is stochastic, occurring base on spontaneous ice nucleation which does not always occur. This may be explained by the nucleation by substances other than water such as macromolecules or dust, termed ‘heterogenous nucleation’(Kanji, Ladino et al. 2017, Shardt, Isenrich et al. 2022). Another possibility is that the inconsistency in the detection of freezing in the midvein may be related to the already high transparency of the leaf venation in optical images making it difficult to detect the subtle change in the reflective index when water freezes.

A notable factor that we did not test in this study is the speed of freezing and thawing. We opted for a slow freeze-thaw trajectory which mimics field conditions but did not test different speeds of freezing or thawing. Slower freezes are thought to increase the likelihood of freeze-thaw embolism (Sevanto, Holbrook et al. 2012) but slower thawing is thought the decrease the loss of conductivity in the xylem (Langan, Ewers et al. 1997). Given the large increase in percentage frozen area between −4°C and −5°C degrees it is possible that that rapidity of freezing over this one degree temperature range precluded embolism formation and that a slower freeze or thaw may have produced different results.

### Freeze-thaw vs. drought induced xylem embolism

Differences in the mechanism behind embolism induced by drought vs. freeze-thaw embolism in the leaf lamina likely explains the contrasting extent of embolism observed in *L. tulipifera* leaves exposed to freezing and drought. The vulnerability of xylem conduits to air-seeding under drought conditions has been linked to anatomical characteristics such as the thickness of the pit membranes in the xylem wall (the site at which air enters, causing embolism due to drought;Thonglim, Delzon et al. 2021, Thonglim, Bortolami et al. 2023) and connectivity of the xylem network (Brodersen and Roddy 2016, Johnson, Brodersen et al. 2020, Mrad, Johnson et al. 2020). Combined, these anatomical features likely explain the pattern of drought-induced embolism we observed in young *L. tulipifera* leaves whereby embolism progressed through the major vein orders terminating in the higher order veins, a pattern which is consistently observed across the leaves of woody and herbaceous angiosperms (Johnson, Jordan et al. 2018, Tonet, Carins-Murphy et al. 2023). In contrast, freeze-thaw embolism, likely caused by gas segregation and subsequent bubble expansion (Sevanto, Holbrook et al. 2012), is thought to be strongly influenced by xylem conduit diameter that determines the size of the resulting bubbles (Pittermann and Sperry 2006).

### What kills *L.* tulipifera leaves during a freeze?

The freeze-induced embolism and damage to living cells detected here highlights that freezing can damage both the water transport and photosynthetic systems in young *L. tulipifera* leaves. It should also be noted however, that regardless of whether embolism occurred, all leaves which showed evidence of mesophyll freezing subsequently died. Cell damage and death was evident in all leaves exposed to temperatures below −4°C, clearly evident in leaf redness immediately following thawing and then subsequent tissue browning in the following days (Fig. S7). This lethal damage was also evident in the depression of chlorophyll fluorescence (Fig. 7). Additionally, *L. tulipifera* leaves presented ‘wilted and wet’ after thawing, a state described by Burke and Gusta et al (1976) in observations of young spring leaves after frost events, attributed to the loss of semi permeability in the cell membranes. The absence of embolism in the lamina of leaves frozen to −5°C indicates that embolism in the lamina is not necessary to induce leaf-death.

## Conclusions

Freezing invariably killed *L. tulipifera* leaves. Death of living tissue was a constant, sometimes accompanied by freeze-thaw embolism which occurred rapidly and was confined to the veins with the largest conduits, in stark contrast to drought-induced embolism. This supports research in stems showing that conduit diameter drives freeze-thaw embolism vulnerability. The progression of freezing through the mesophyll in patches bounded by tertiary veins implicates BSEs in driving lateral ice propagation. While freezing in the mesophyll conclusively led to death in *L. tulipifera* leaves, a significant proportion of xylem vessels were inherently vulnerable to freeze-thaw embolism, showing that spring frost results in a dual mode of lethality in *L. tulipifera* leaves.

## Supplementary Data

**Figure S1:** Transverse light microscope section of the lamina of a *Liriodendron tulipifera* leaf highlighting a bundle-sheath extension.

**Figure S2:** Comparison of natural and experimental freezing trajectories.

**Video 1:** The pattern of freezing and thawing detected in a leaf frozen to −5 ° C with time-lapse imaging.

**Figure S3:** Examples of the ‘patchy’ nature of freezing observed in *Liriodendron tulipifera* leaves frozen to air temperatures of both −4 ° C and −5 ° C.

**Figure S4:** Chlorophyll fluorescence (Fv/Fm) in *Liriodendron tulipifera* leaves which were placed in the freezer but did not show visible signs of freezing.

**Figure S5:** Comparison of optically resolved freezing and cell damage shown through imaging fluorescence in *Liriodendron tulipifera leaves*.

**Figure S6:** The total embolism overlayed onto initial images of each the six *Liriodendron tulipifera* leaves in which embolism was observed.

**Video 2**: High resolution freezing in a *Liriodendron tulipifera* leaf.

**Video 3**: High resolution GIF of thawing of a *Liriodendron tulipifera* leaf.

**Figure S7:** *Liriodendron tulipifera* trees shortly after they were exposed to −4°C or −5°C freeze treatments.

## Acknowledgements

We would like to thank Peter Johnson for his assistance in designing the ‘Envirologger’ monitoring systems.

## Author contributions

KMJ and CRB conceived and designed the study. KMJ and CRB conducted experiments and measurements using optical cameras and microscopy, while KMJ and MS designed and conducted the collection of high-resolution imagery with guidance from DG, RWS and ERD. KMJ and CRB wrote the manuscript with contributions from all authors.

## Data availability

The data will be made available by the corresponding authors upon reasonable request.

