## Supplementary data for "Ice and air: Visualisation of freeze-thaw embolism and freezing spread in young *L. tulipifera* leaves"

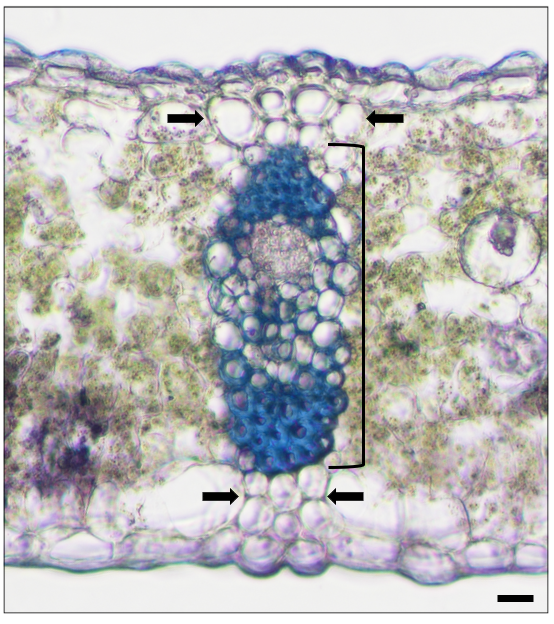

**Figure S1:** Transverse light microscope section of the lamina of a *Liriodendron tulipifera* leaf highlighting a bundle-sheath extension. This image includes a third order vein (black bracket) and bundle sheath extension (denoted by black arrows) extending to the upper and lower leaf surfaces. The black scale bar in the bottom right is 2 microns.

**Figure S2:** Comparison of natural and experimental freezing trajectories. Freezing trajectories from a weather station at Yale Myers Forest where *Liriodendron tulipifera* naturally occurs, taken at 3 minute
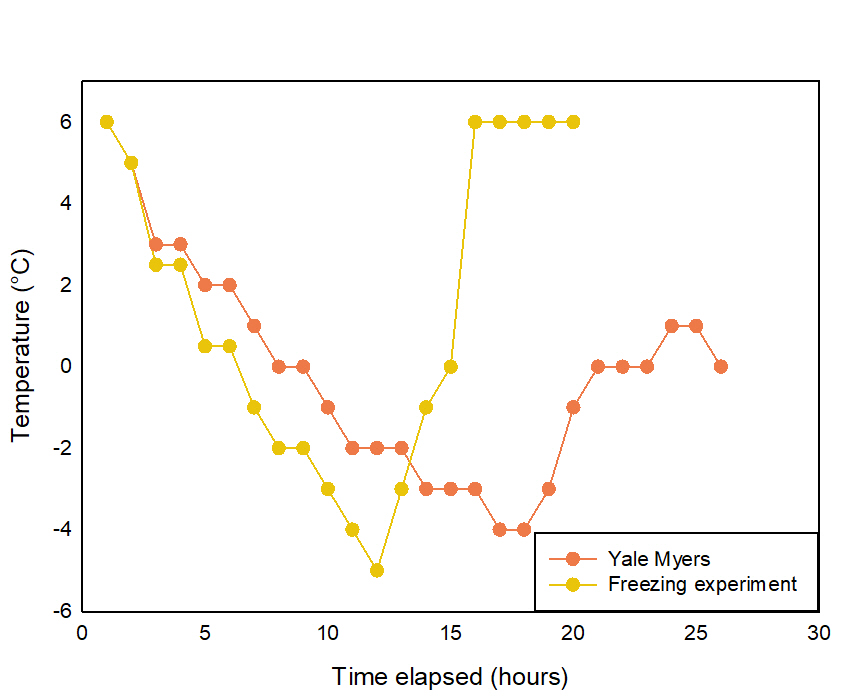
intervals and averaged to 1 hour temperature readings (red) and the temperatures that *L. tulipifera* trees were exposed to in freezers (yellow)

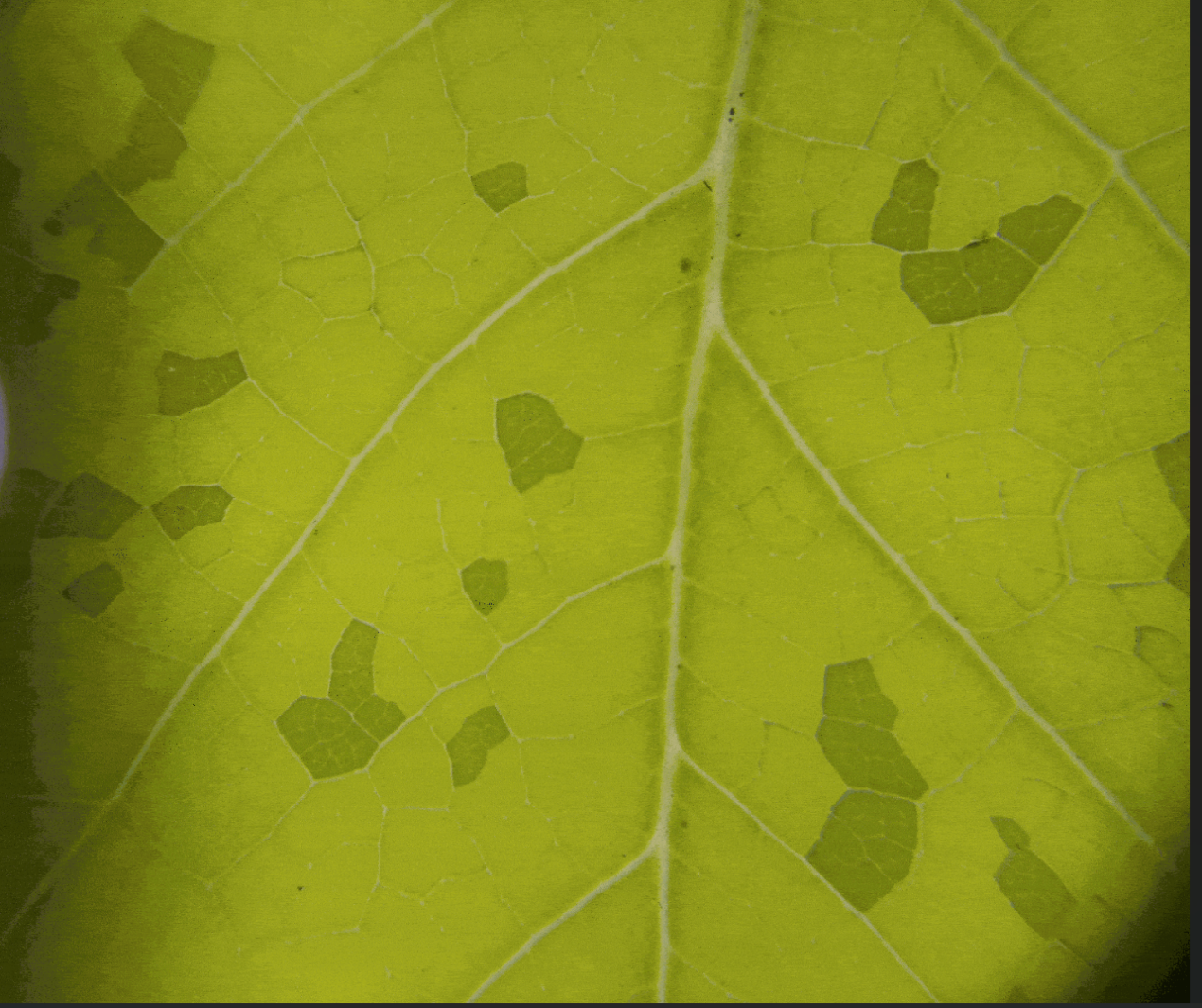

**Video 1:** [PLACEHOLDER IMAGE] The pattern of freezing and thawing detected in a leaf frozen to -5 ° C with time-lapse imaging.

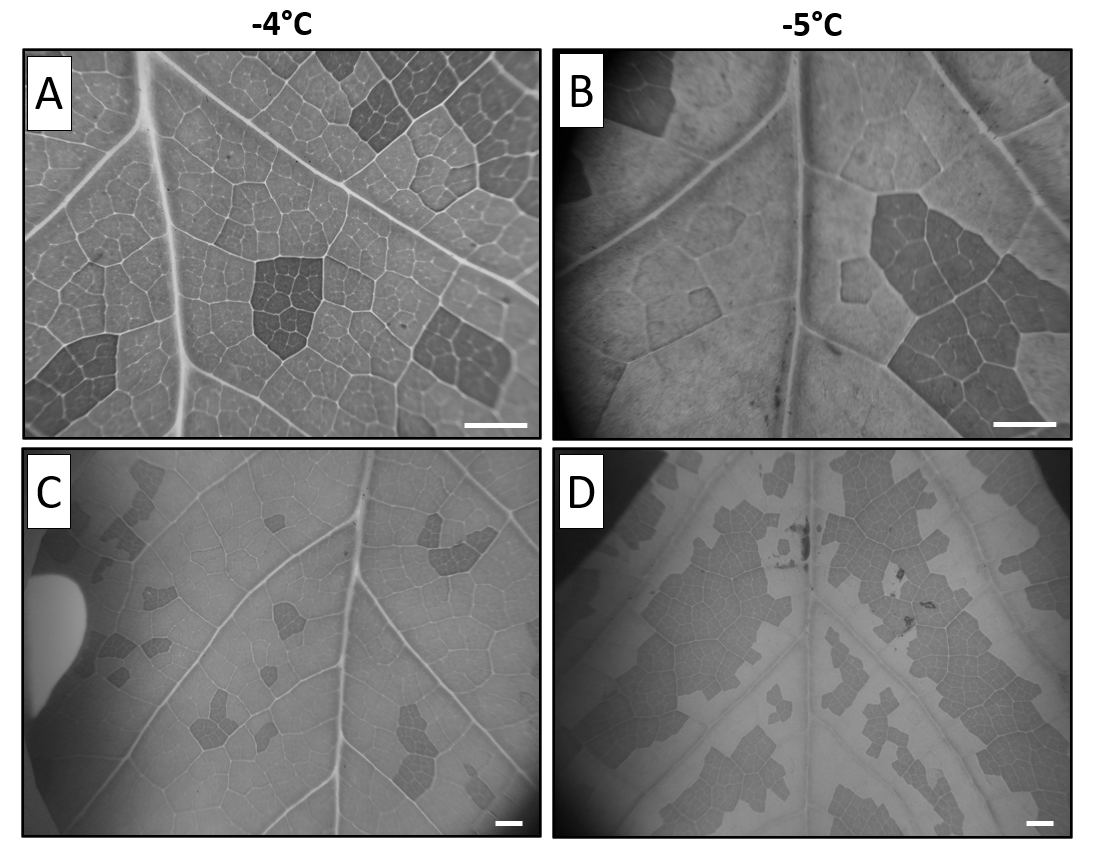
**Figure S3:** Examples of the ‘patchy’ nature of freezing observed in *Liriodendron tulipifera* leaves frozen to air temperatures of both -4 ° C and -5 ° C. Shown at magnification (A and B) and lower magnification (C and D). The white scale bars are 2mm**.**

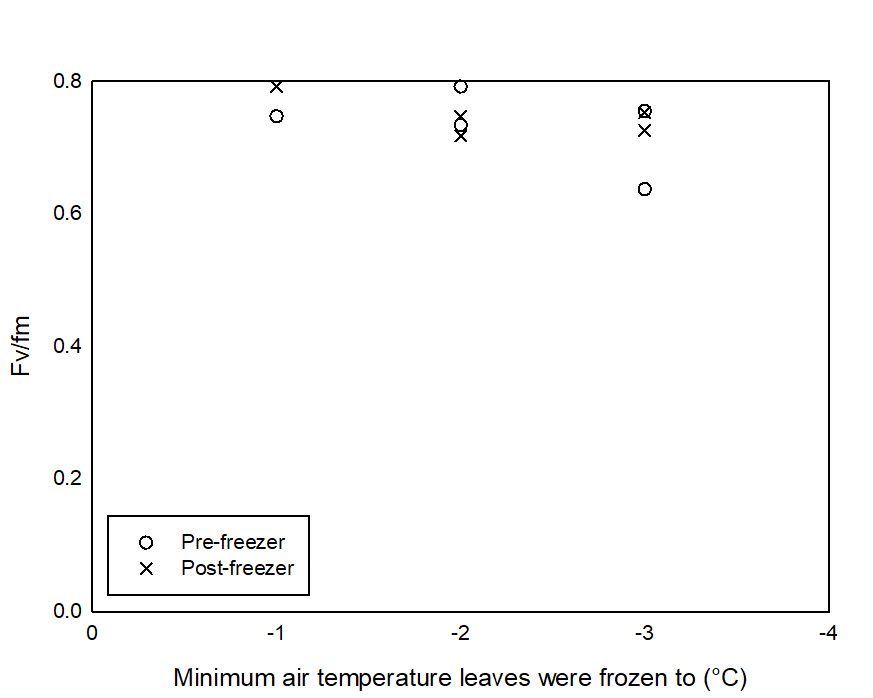

**Figure S4:** Chlorophyll fluorescence (Fv/Fm) in *Liriodendron tulipifera* leaves which were placed in the freezer but did not show visible signs of freezing. Pre-freezer measurements are denoted by circles while post freezer measurements are denoted by crosses and plotted against the minimum air temperature leaves were frozen to.

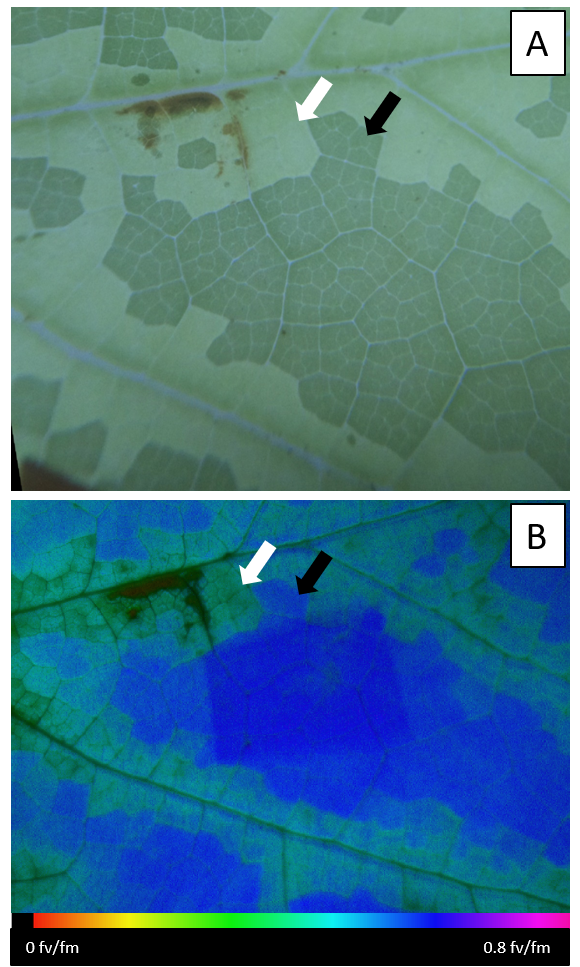

**Figure S5:** Comparison of optically resolved freezing and cell damage shown through imaging fluorescence in *Liriodendron tulipifera leaves*. The patchy pattern of freezing (a) and consequent depressed fluorescence (b) after freezing along the midrib and secondary veins in an *L. tulipifera* leaf. The colour scale at the bottom represents the spectrum of fluorescence from and fv/fm of 0 (dead cells) to 0.8 (healthy cells). The white and black arrows indicate patches of frozen and unfrozen tissue respectively in both (a) and (b).

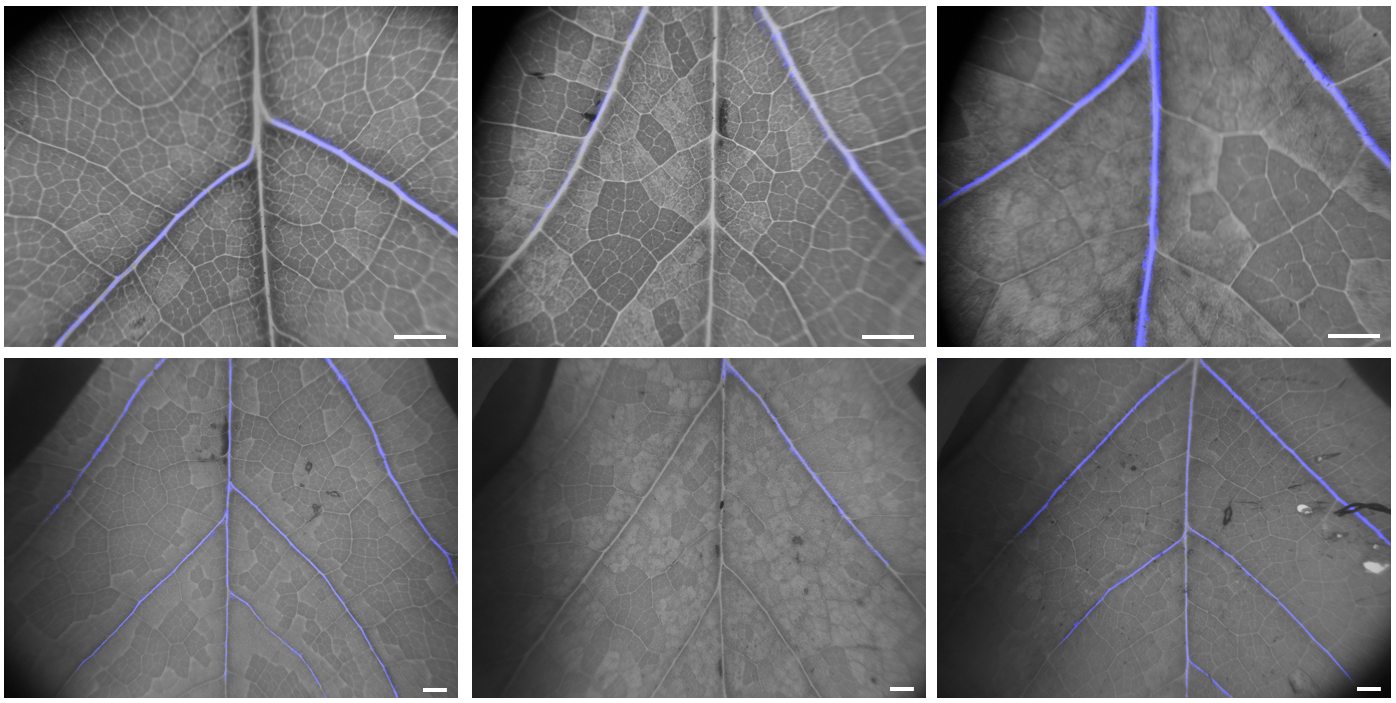

**Figure S6:** The total embolism overlayed onto initial images of each the six *Liriodendron tulipifera* leaves in which embolism was observed. Embolism is shown in blue while the leaves are grey. The white scale bars in the bottom right rand corner of the images are 2mm.

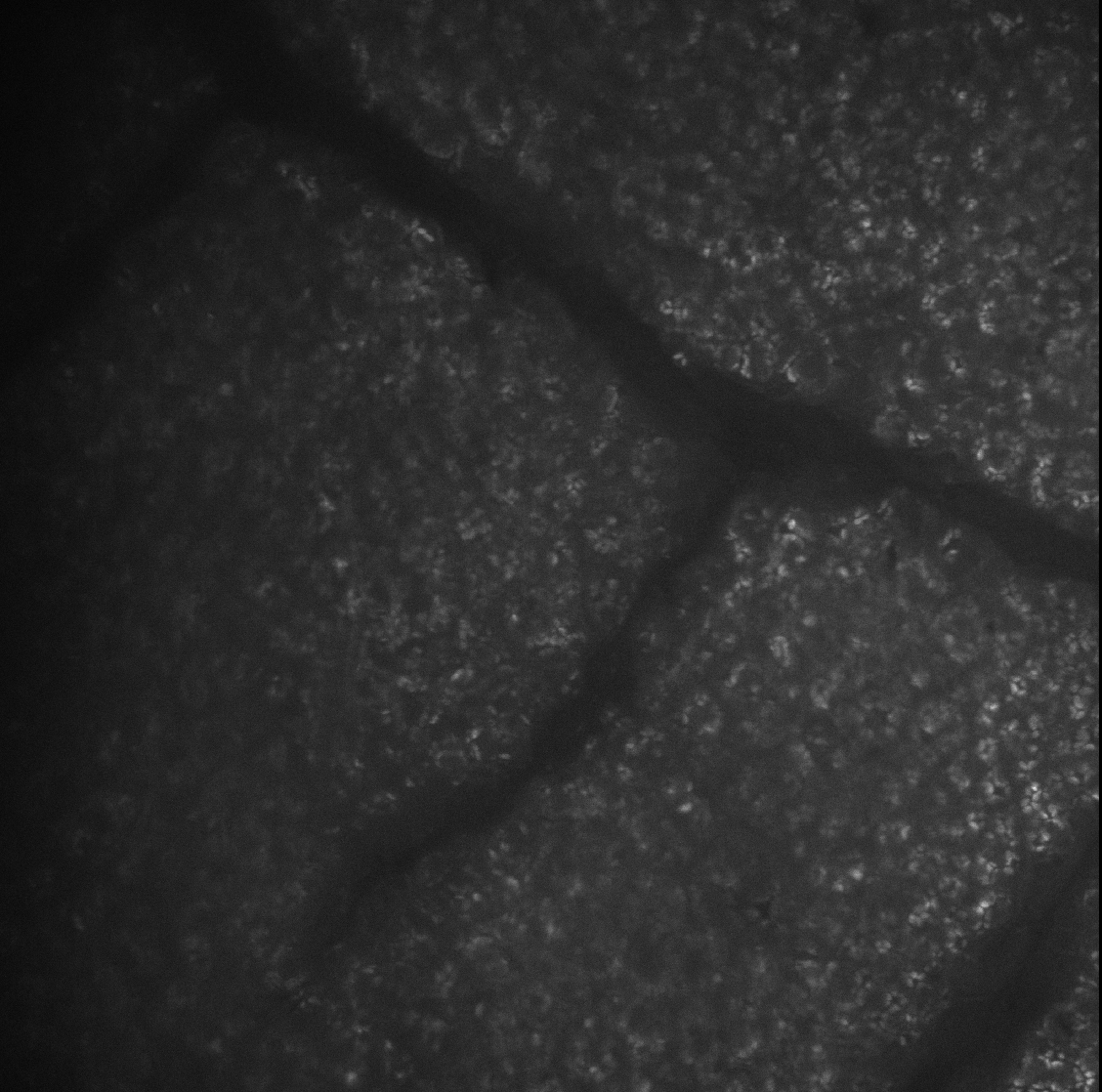

**Video 2**: [PLACEHOLDER IMAGE] High resolution freezing in a *Liriodendron tulipifera* leaf. The freezing front initiates in the bottom Lefthand corner and propagates towards the middle of the field of view before reaching a third order vein boundary. The freezing front then continues from the bottom righthand corner towards the middle.

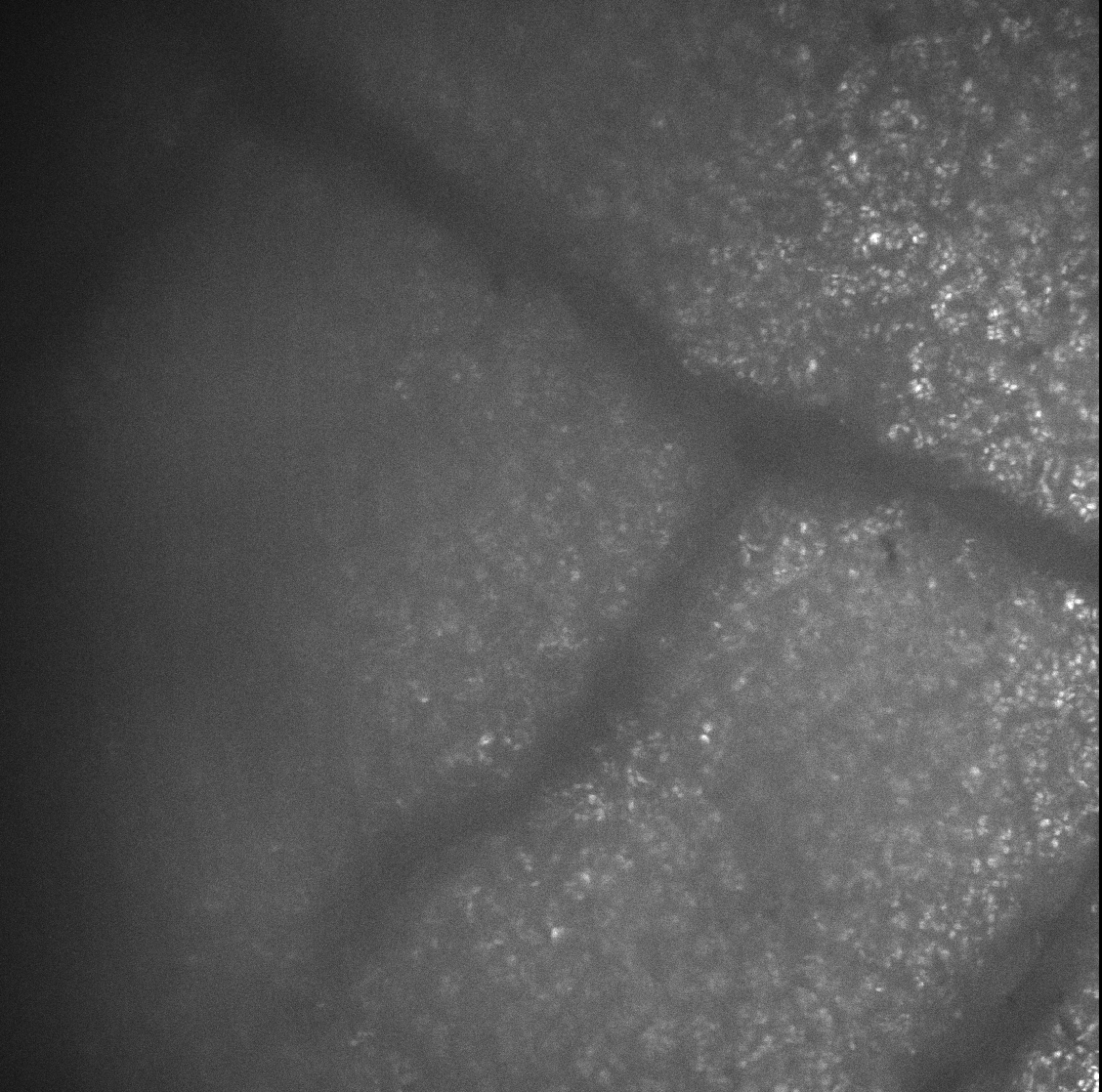

**Video 3**: [PLACEHOLDER IMAGE] High resolution thawing of a *Liriodendron tulipifera* leaf. Thawing appears to occur in a generally uniform manner, with slight directionality (in the opposite direction of freezing, Video 1) from the top right to the bottom left.

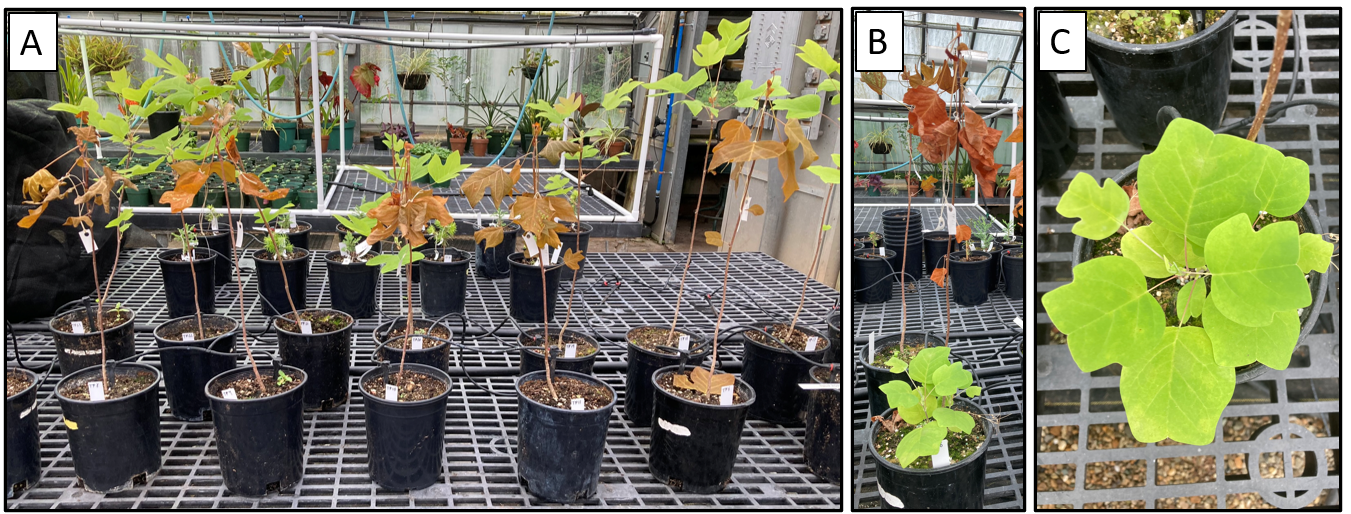

**Figure S7:** *Liriodendron tulipifera* trees shortly after they were exposed to -4°C or -5°C freeze treatments. The front row shows trees exposed to these temperatures, with browning evident in 100% of their leaves (A). Panels (B) and (C) show resprouting at the base of a tree. This occurred in all but one trees which was exposed to -4°C and -5°C freezes in the weeks following the freeze treatment.
